# Extremely bright, near-IR emitting spontaneously blinking fluorophores enable ratiometric multicolor nanoscopy in live cells

**DOI:** 10.1101/2021.06.02.446776

**Authors:** Jonathan Tyson, Kevin Hu, Shuai Zheng, Phylicia Kidd, Neville Dadina, Ling Chu, Derek Toomre, Joerg Bewersdorf, Alanna Schepartz

**Affiliations:** Department of Chemistry, University of California, Berkeley, USA; Department of Molecular and Cellular Biology, University of California, Berkeley, USA; California Institute for Quantitative Biosciences (QB3), University of California, Berkeley, USA; Department of Cell Biology, Yale School of Medicine, New Haven, CT, USA; Department of Chemistry, Yale University, New Haven, CT, USA; Department of Molecular, Cellular, and Developmental Biology, Yale University, New Haven, CT, USA; Department of Biomedical Engineering, Yale University, New Haven, CT, USA; Kavli Institute for Neuroscience, Yale School of Medicine, New Haven, CT, USA; Nanobiology Institute, Yale University, West Haven, CT, USA

## Abstract

New bright, photostable, emission-orthogonal fluorophores that blink without toxic additives are needed to enable multi-color, live-cell, single-molecule localization microscopy (SMLM), especially for experiments that demand ultra-high-resolution live imaging. Here we report the design, synthesis, and biological evaluation of Yale_676sb_, a photostable, near-IR emitting fluorophore that achieves these goals in the context of an exceptional quantum yield (0.59). When used alongside HMSiR, Yale_676sb_ enables simultaneous, live-cell, two-color SMLM of two intracellular organelles (ER + mitochondria) with only a single laser and no chemical additives.

---

Single-molecule localization microscopy (SMLM)^1–4^ is a powerful technique for visualizing intracellular architecture at the nanoscale^5^ and across large fields of view (FOV).^6^ The technique is characterized by the detection and localization of fluorescent markers that cycle rapidly between emissive (ON) and non-emissive (OFF) states. For optimal results, the sample and imaging conditions must maintain the majority of fluorescent markers in the OFF state, such that the neighboring molecules in the emissive ON state can be treated as sparse single emitters.^7–11^ Organic fluorophores are favored over fluorescent proteins for SMLM because they are generally brighter, more photostable, and because their photophysical properties can be finetuned using chemistry.^7,12–17,17–21^ The challenge is that many SMLM-compatible organic fluorophores require the addition of exogenous nucleophiles, redox modulators, and/or oxygen depletion systems to switch efficiently between ON and OFF states. These additives can be cytotoxic and damage or alter biological samples.^7,12,13,15,17^ An additional challenge is that many established SMLM-compatible fluorophores are cell-impermeant^7,12,13,20,22,23^ and/or require cytotoxic high-power and/or short-wavelength lasers.^7,12,16,18,22,24,25^

The spontaneously blinking fluorophore (SBF) hydroxymethyl Si-rhodamine (HMSiR)^15^ (Figure 1A) reported by Urano and coworkers overcomes many of these limitations. It is cell-permeant, photostable, and has been proposed to cycle rapidly between ON and OFF states by virtue of a pH-dependent spirocyclization reaction that occurs in the absence of chemical additives^15^ (Figure 1A). For HMSiR, the midpoint of this pH-dependent equilibrium (referred to as p*K*_cycl_) occurs at approximately pH 6.0. Thus, at pH 7.4 roughly 98% of the HMSiR molecules in solution occupy the OFF state, which enables facile detection and localization of the sparse subset of molecules that are emissive (ON).^15^ HMSiR’s cell permeability, photostability, and ability to blink in the absence of chemical additives has enabled multiple minimally invasive single-color SMLM experiments, including those that visualize organelle membrane dynamics in live cells for extended times,^26^ others that resolve the morphology of dopaminergic neurons in an intact *Drosophila melanogaster* adult brain,^27^ and still others that enable turn-on visualization of intracellular protein targets.^28^

**Legend for Figure 1.**
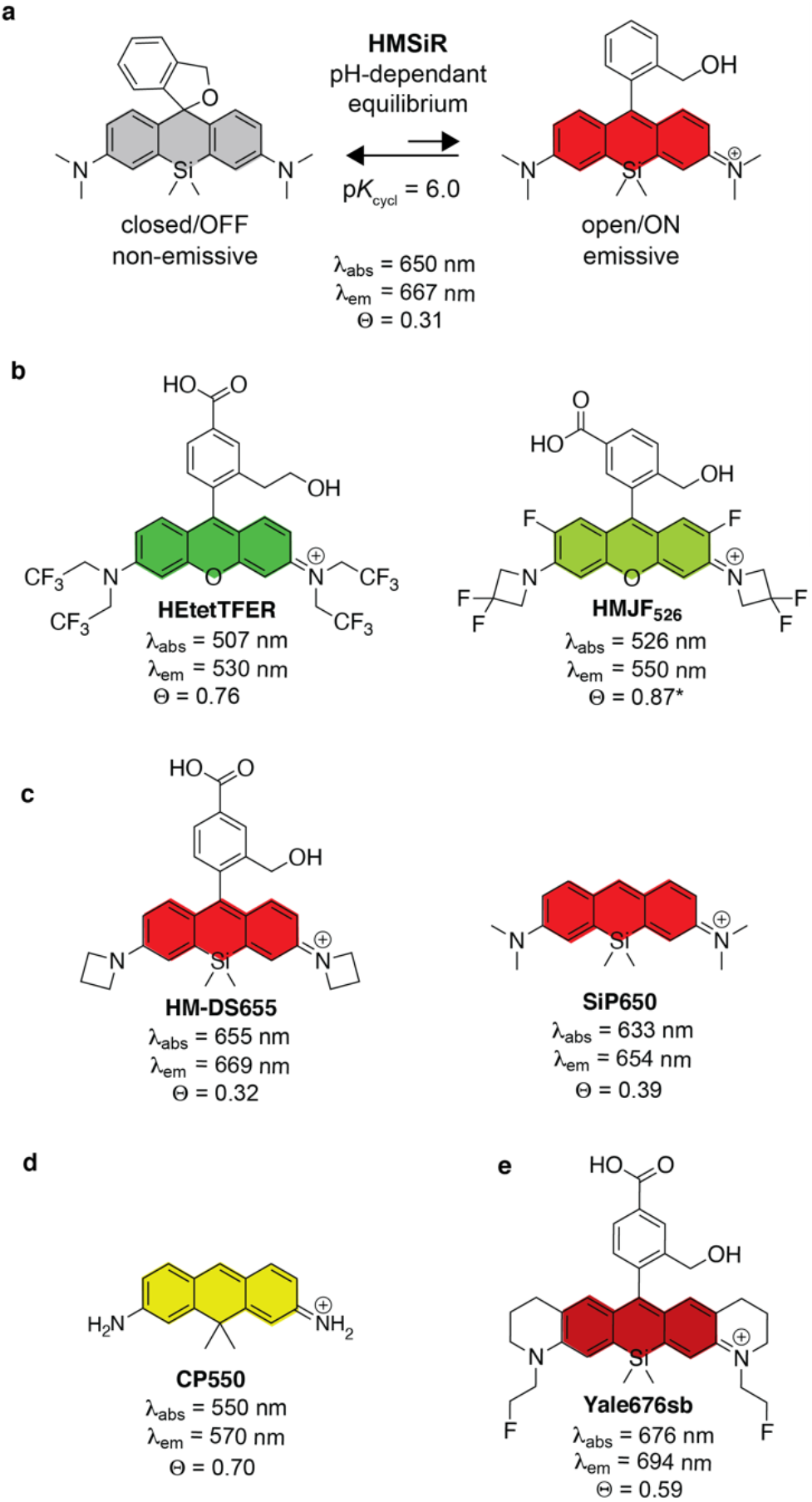
(A) Structure and pH-dependent equilibrium of the spontaneously blinking fluorophore HMSiR.^15^ (B-D) Structures of previously reported fluorophores considered as potential HMSiR partners for multicolor live-cell SMLM^16,18,21,29^ (E) Structure of the spontaneously blinking fluorophore reported herein, Yale_676sb_.

Despite these advances, there remains a need for new SBFs that effectively partner with HMSiR to enable multicolor live-cell SMLM experiments without the need for chemical additives or photoactivation^13^. Although two green-emitting SBFs whose emission spectra are separable from HMSiR have been reported (Figure 1B),^18,19^ including one (HEtetTFER) that can be paired with HMSiR for two-color SMLM in fixed cells,^18^ their use demands high-intensity lasers that excite at 488 and 561 nm, respectively, which can induce substantial cytotoxicity as phototoxicity is especially pronounced in the blue and green spectrum.^14^ Two other previously reported SBFs are excitable in the far-red/near-IR (Figure 1C),^20,21^ but they are spectrally indistinguishable from HMSiR and are therefore not suitable for two-color experiments. Although both the fluorescent protein mEos3.2 as well as CP550 (Figure 1D), a carbopyronin fluorophore that reacts irreversibly with intracellular glutathione^20^ have been paired with HMSiR for two-color live cell SMLM,^15,20^ these experiments require an additional ~560 nm laser, which is inferior to red-light excitation for live-cell microscopy.^10,14^ Furthermore, sequential multi-color imaging with multiple lasers is slow, making it more prone to sample motion artifacts. Finally, the spontaneously blinking carborhodamine, HMCR550, which was designed using quantum calculations, has an excitation maximum at 560 nm and would likewise require multiple lasers to pair with HMSiR (650 nm excitation) for a two-color live cell SMLM experiment.

Here we report the rational design of a new near-IR emitting SBF that pairs effectively with HMSiR to enable simplified two-color SMLM experiments in live cells (Figure 1D). Yale_676sb_ emits at 694 nm, the longest wavelength of any reported SBF, and possesses, to our knowledge, a higher quantum yield (0.59) than any previously reported nanoscopy-compatible Si-rhodamine (SiR) fluorophore. Yale_676sb_ and HMSiR can be excited simultaneously with a single 642-nm laser and imaged ratiometrically for simultaneous multicolor SMLM of two distinct intracellular organelles (ER + mitochondria) in live cells.

## Results

### New spontaneously blinking fluorophores: Design considerations

Three distinct chemical and photophysical properties are needed to ensure compatibility with HMSiR for ratiometric two-color, live-cell SMLM. The first is an emission maximum >690 nm to ensure adequate separation from HMSiR (emission maximum = 670 nm) *via* ratiometric imaging.^5,30,31^ The second is a p*K*_cycl_ value between 5.3 and 6.0 to ensure the sparsity of emissive/ON molecules.^15,21^ The third requirement is a high quantum yield; although a quantum yield >0.2 can yield respectable SMLM images, higher values are always more desirable.^13,22^ The challenge is that the quantum yields of rhodamine-based fluorophores typically decrease as the absorption and emission maxima increase (Supplemental Figure 1). As a result, molecules that absorb and emit at higher, less cytotoxic wavelengths that are compatible with live-cells are relatively dim. This correlation is reflected in the relatively low quantum yield of HMSiR (0.31) when compared to those of the green-light emitting SBFs HMJF526 (0.87)^16^ and HEtetTFER (0.76)^18^. We therefore sought a design approach that would yield fluorophores possessing both long-wavelength emission and high quantum yield.

### HMSiR_indol_, HMSiR_julol_, and HMSiR_THQ_

Previous work has demonstrated that introduction of heterocyclic indoline^32,33^, julolidine^34^, or tetrahydroquinoline^32^ moieties into the core of a SiR chromophore can shift the excitation and emission maxima by up to 50 nm relative to SiR itself (Figure 2A). To evaluate whether these effects would be preserved in the context of a HMSiR core, we synthesized HMSiR, as well as the heterocyclic derivatives HMSiR_indol_, HMSiR_julol_, and HMSiR_THQ_ (Figure 2B and Supplementary Schemes 1-4) according to a recently reported general method for Si-rhodamine fluorophore synthesis.^35^ We then characterized the photophysical properties and aqueous spirocyclization equilibrium (p*K*_cycl_) of each new fluorophore (Figure 2B-D).

**Legend for Figure 2.**
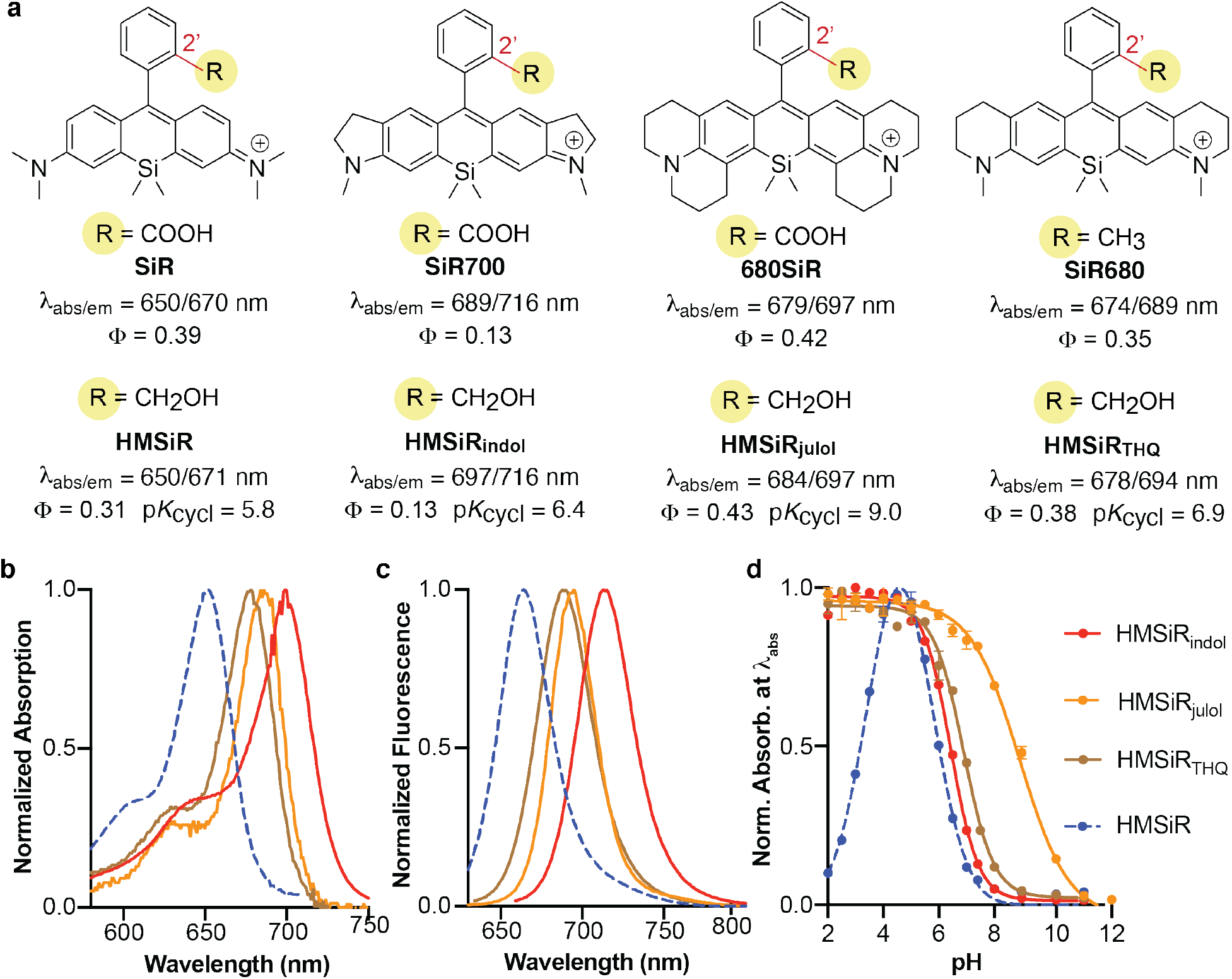
(A) Structures and photophysical properties (ƛ_abs_, ƛ_em_, and ϕ) of previously reported SiR fluorophores with red-shifted absorption and emission spectra and HMSiR analogs HMSiR, HMSiR_indol_, HMSiR_julol_, and HMSiR_THQ_. Normalized (B) absorption and (C) emission spectra of HMSiR, HMSiR_indol_, HMSiR_julol_, and HMSiR_THQ_ in 0.2 M sodium phosphate (pH = 4.5 for HMSiR, pH = 2.0 for HMSiR_indol_, HMSiR_julol_ and HMSiR_THQ_). (D) pH-dependent change in absorbance of 2 μM HMSIR (650 nm), HMSiR_indol_ (697 nm), HMSiR_julol_ (684 nm), and HMSiR_THQ_ (678 nm) as a function of pH in 0.2 M sodium phosphate buffer at room temperature. The absorbance of each fluorophore was monitored at the wavelength of maximal absorbance in (B).

Each of the new fluorophores displayed absorption (Figure 2B) and emission (Figure 2C) maxima that were red-shifted by at least 23 nm relative to HMSiR, with the emission maxima increasing in the order HMSiR < HMSiR_THQ_ < HMSiR_julol_ < HMSiR_indol_. As expected, the absorption and emission maxima of the HMSiR series were nearly identical to those of the analogous SiR variants reported previously.^32–34^ The p*K*_cycl_ of each new HMSiR analog was determined from a plot of the pH-dependence of the absorption of each fluorophore at the absorption maximum of the open/ON form (Figure 2D and Supplementary Figure 2); the p*K*_cycl_ is the pH at which the concentration of the open/ON state equals that of the closed/OFF state.^15^

The p*K*_cycl_ values of HMSiR_indol_ and HMSiR_THQ_ were 6.4 and 6.9, respectively, both significantly higher than the value for HMSiR (6.0). The p*K*_cycl_ value of HMS_julol_ (p*K*_cycl_ = 9.0) was shifted even more dramatically, presumably because the additional electron-donating alkyl groups disfavor cyclization. A related previously reported rhodamine analog with julolidine groups also displayed a high p*K*_cycl_. value.^15^ The absorbance *versus* pH curves for HMSiR_indol_, HMSiR_julol_, and HMSiR_THQ_ are sigmoidal, whereas that of HMSiR is bell-shaped due to cyclization of the protonated fluorophore at low pH; this protonation is disfavored when the exocyclic amine is constrained by a 5- or 6-membered ring.^15,36^

The final criterion needed to ensure compatibility with HMSiR for ratiometric two-color, live-cell SMLM is a high quantum yield. The quantum yields measured for HMSiR_indol_, HMSiR_julol_, and HMSiR_THQ_ also paralleled the values for the analogous SiR variants; the quantum yield of HMSiR_indol_, like SiR700, was low (0.13), whereas those of HMSiR_julol_ and HMSiR_THQ_ (0.43 and 0.38, respectively) were comparable to that of HMSiR (0.31) (Supplementary Figure 3).

These data indicate that neither HMSiR_THQ_, HMSiR_julol_, nor HMSiR_indol_ possess the characteristics necessary to partner with HMSiR for two-color SMS nanoscopy. Although all three fluorophores exhibit emission maxima that are shifted by at least 23 nm from that of HMSiR, and HMSiR_julol_ and HMSiR_THQ_ display acceptable quantum yields (0.43 and 0.38), none feature p*K*_cycl_ values low enough to prevent significant multi-emitter artifacts at physiological pH. In each case, chemical modifications are needed to increase the electrophilicity of the xanthene core, favor spirocyclization, and decrease p*K*_cycl_. Ideally, these modifications should also increase quantum yield to increase brightness and resolution, but as outlined below, this goal is complicated by the complex interplay between emission maximum, p*K*_cycl_ and quantum yield.

### Interplay between emission maximum, p*K*_cycl_ and quantum yield

The quantum yields of rhodamine fluorophores are limited by a non-radiative decay process known as twisted intramolecular charge transfer (TICT)^37–39^. TICT involves the excited-state transfer of an electron from the exocyclic nitrogen of the fluorophore to the neighboring carbon pi system with concomitant twisting of the Caryl-N bond; the charge-separated state subsequently decays to the ground state without emission of a photon. Processes that decrease the Caryl-N bond rotation increase the quantum yield. For example, the quantum yields of rhodamine B and tetramethyl rhodamine (TMR) are higher in viscous solvents^37^ and at low temperature where C_aryl_-N bond rotation is inhibited^37,40^. Indeed, the modestly increased quantum yields of HMSiR_julol_ (0.43) and HMSiR_THQ_ (0.38) relative to HMSiR (0.31) can also be ascribed to restricted C_aryl_-N bond rotation^34^, although these effects appear to be less dramatic in the SiR series than with conventional rhodamines: Rhodamine 101, the rhodamine analog of HMSiR_julol_, displays a near-perfect quantum yield of 0.99.^40^

TICT is also inhibited in fluorophores in which the ionization potential (IP) of the exocyclic nitrogen is increased by electron-withdrawing groups (EWGs).^18,38,41^ Addition of EWGs to a fluorophore core also decreases p*K*_cycl_ by lowering the energy of the fluorophore’s lowest unoccupied molecular orbital (LUMO).^15,18^However, addition of EWGs typically induces moderate to large decreases in excitation and emission wavelength maxima. For example, an EWG-containing fluorophore reported by Lv *et al.* possesses an exceptional quantum yield (0.66) but is blue-shifted by ~20 nm relative to HMSiR (ƛ_abs_/ƛ_em_ = 631/654 nm).^41^ We hypothesized that combining the effects of restricted aryl-N bond rotation with an EWG would simultaneously reduce p*K*_cycl_ and increase quantum yield by inhibiting TICT. If these changes were introduced into the HMSiR_THQ_ scaffold, even a moderate decrease in excitation and emission maxima would not jeopardize the emission shift needed to remain orthogonal to HMSiR. HMSiR_THQ_ was preferred as a starting point because its p*K*_cycl_ (6.9) and quantum yield (0.38) are both close to that of HMSiR, in contrast to HMSiR_indol_, whose quantum yield is low (0.13) or HMSiR_julol_, whose p*K*_cycl_ is very high (9.0).

### Design of the bright, near-IR emitting SBF, Yale_676sb_

To test this hypothesis, we synthesized Yale_676sb_, a variant of HMSiR_THQ_ in which two N-methyl groups were replaced symmetrically by mono-fluorinated N-ethyl groups (Figure 3 and Supplementary Scheme 5). As predicted, Yale_676sb_ was characterized by a 10-fold more favorable spirocyclization equilibrium than HMSiR_THQ_ (p*K*_cycl_ = 5.9 *vs.* 6.9) and a greatly improved quantum yield (0.59 *vs.* 0.38) (Supplementary Figure 3). Interestingly, Yale_676sb_ exhibited absorption and emission ƛ_max_ that are both virtually identical to those of HMSiR_THQ_. Addition of a stronger di-fluorinated N-ethyl group to generate Cal664sb resulted in a further increase in quantum yield to 0.74 (Supplementary Figure 3), but in this case led to an emission ƛ_max_ that was too close to that of HMSiR (667 nm *vs.* 677 nm) to support two-color ratiometric imaging. These photophysical properties associated with Yale_676sb_ suggest that it should be an ideal partner for HMSiR: an emission maximum >690 nm, a p*K*_cycl_ value between 5.3 and 6.0, and a high quantum yield. The quantum yield of Yale_676sb_ (0.59) is, to our knowledge, higher than any Si-rhodamine derivative prepared and utilized for fluorescence nanoscopy.

**Legend for Figure 3.**
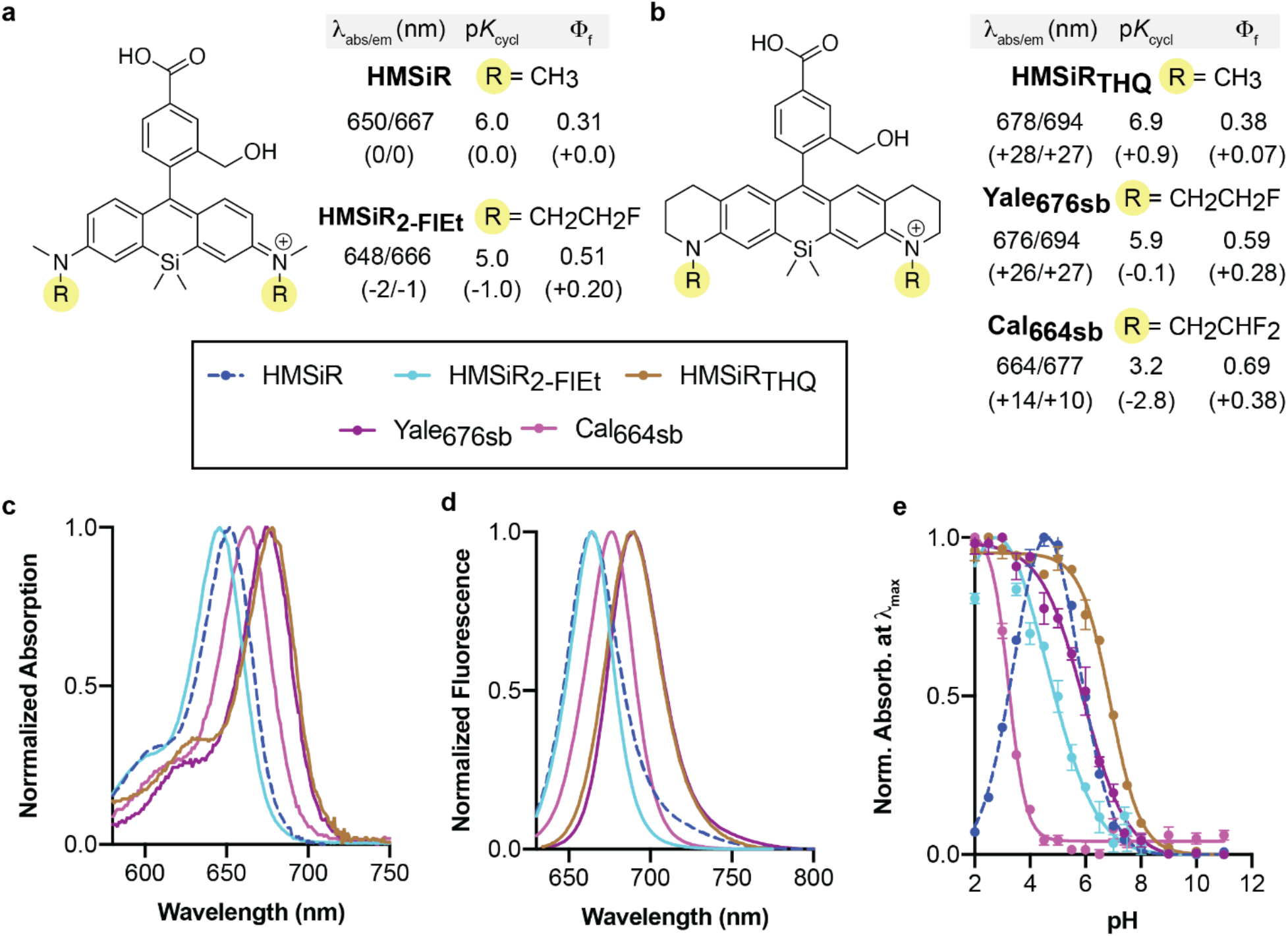
Structures and photophysical properties (ƛ_abs_, ƛ_em_, and ϕ) of (A) HMSiR, HMSiR2FlEt, (B) HMSiR_THQ_, Yale_676sb_ and Cal664sb. Normalized (C) absorption and (D) emission spectra of HMSiR, HMSiR_indol_, HMSiR_julol_, and HMSiR_THQ_ in 0.2 M sodium phosphate (pH = 4.5 for HMSiR, pH = 2.0 for HMSiR_indol_, HMSiR_julol_ and HMSiR_THQ_). (E) pH-dependent spirocyclization equilibria. Normalized absorption of open form of 2 μM HMSiR, HMSiR_2-FlEt_, HMSiR_THQ_, Yale_676sb_ and Cal664sb as a function of pH in 0.2 M sodium phosphate buffer at room temperature.

To deconvolute the effects of aryl-N bond rotation and the mono-fluoro electron withdrawing group, we also prepared HMSiR_2-FlEt_, which carries the same mono-fluorinated N-ethyl groups but allows aryl-N bond rotation (Supplementary Scheme S7). HMSiR_2-FlEt_ was characterized by a minimal change in absorption and emission ƛ_max_ relative to HMSiR; however, it displayed a 10-fold more favorable spirocyclization equilibrium than HMSiR (p*K*_cycl_ = 5.0 *vs.* 6.0), a value too low for efficient blinking at physiological pH of 7.4.^21^ Its improvement in quantum yield was more modest relative to Yale_676sb_ (0.51 vs 0.59). These comparisons emphasize the benefits of combining restricted aryl-N bond rotation with an EWG.

### Evaluation of the single molecule properties of Yale_676sb_

To ensure that the bulk photophysical parameters of Yale_676sb_ would translate into efficient single-molecule parameters, we evaluated its properties under SMLM imaging conditions. We quantified the ‘on-time’ to determine the dye’s compatibility with HMSiR by imaging single dye molecules immobilized on glass coverslips (Supplementary Figure 4). By monitoring individual molecules, we were able to determine the on-time to evaluate the compatibility of Yale_676sb_ and HMSiR. Because both dyes are imaged on the same camera using ratiometric imaging, similar on-times allow a single camera integration time to acquire data effectively from both fluorophores.^42^ From these data, we determined that Yale_676sb_ has an on-time of 4.5 ms at pH 7.4, which is close to the ~10 ms on-time reported for HMSiR, and in theory should allow even faster imaging.^15^ This short on-time, combined with the high quantum yield, also makes the Yale_676sb_/HMSiR combination suitable for high-speed imaging, with camera frame rates as high as 400 frames per second (fps). With an off-time of 3.8 s, we expect an on-fraction or duty cycle of 0.0012.

### Single-color live-cell SMLM with Yale_676sb_

We next tested whether Yale_676sb_ would support single-color, live-cell SMLM imaging. U2-OS cells that were engineered to overexpress the endoplasmic reticulum (ER)-localized protein Halo-Sec61β^43^ were treated with 300 nM Yale_676sb_-CA (Supplementary Scheme S8) for 30 min, then washed and immersed in a standard live-cell imaging solution using a custom-built SMLM instrument (see Supplementary Methods). Figure 4A shows a representative super-resolution image (out of n = 16 images) acquired over 5 seconds. These images revealed multiple tubules in the cell periphery that were ~99 ± 15 nm (mean ± s.d.) wide, a value comparable to ER morphology metrics acquired using both STED and 4Pi-SMS.^44^ A time series illustrates changes in ER morphology that occur over the course of 10 seconds (Figure 4B). On average, we detected ~800 photons per blink, corresponding to a localization precision distribution with a peak at ~20 nm (Figure 4C, D).

**Legend for Figure 4.**
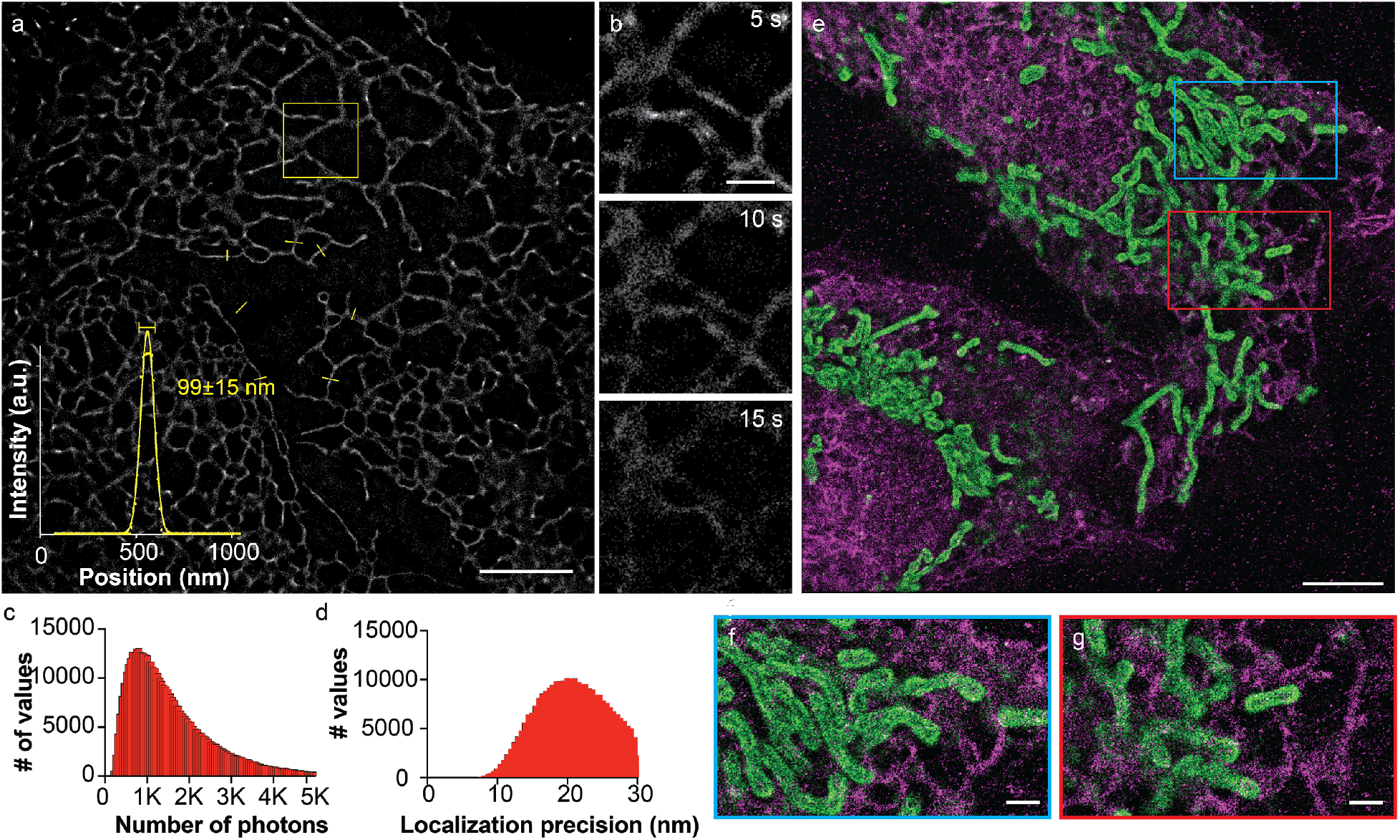
(A) Super-resolution image of the ER in U2-OS cells using Yale_676sb_. Average reconstructed signal as a function of position along the seven line profiles indicated by yellow lines is shown. Inset (B) shows dynamic ER network remodeling. Histograms illustrating the range in the number of photons (C) and localization precision (D) associated with single-molecules in (A). (E) Two-color super-resolution image of the ER and mitochondria in U2-OS cells using Yale_676sb_ (magenta) in conjunction with HMSiR (green). Insets (F-G) show superresolved mitochondrial and ER networks in close proximity. Scale bars 5 μm for (A) and (E), 1 μm for (B) and (F). All reconstructions using 5 s of acquired frames.

### Ratiometric two-color live-cell SMLM with Yale_676sb_ and HMSiR

Next we sought to evaluate whether Yale_676sb_ would support live-cell multicolor imaging in combination with HMSiR. U2-OS cells were transiently transfected with Halo-Sec61β (to reveal the ER) and SNAP-OMP25 (to reveal the outer mitochondrial membrane), treated with Yale_676sb_-CA and HMSiR-BG, and imaged using the identical SMLM setup. As predicted from the absorption and emission maxima of Yale_676sb_ and HMSiR, both dyes could be excited with the same 642-nm laser and ratiometrically separated from two simultaneously acquired images detecting the emission wavelength ranges of 650-680 nm and 680-750 nm, respectively (Supplementary Figure 5). Figure 4E shows a resulting two-color super-resolution image, accumulated over 5 s, revealing the intertwined mitochondrial and ER networks of the cell. We detected comparable average photon numbers per frame for the two dyes (~500 and 590 photons for Yale_676sb_ and HMSiR, respectively), especially given that the filters and excitation wavelength were optimized for HMSiR.

## Conclusions

In summary, here we report a new spontaneously blinking Si-rhodamine, Yale_676sb_ that can be used alongside HMSiR to enable two-color ratiometric SMLM in living cells in physiological media. This new experiment was facilitated by three unique photophysical metrics associated with Yale_676sb_: (1) an exceptionally high quantum yield for a silicon rhodamine derivative (0.59); (2) an unusually long emission maximum (694 nm); and (3) a p*K*_cycl_ value (5.9) that is nearly identical to that of HMSiR (6.0).

The unique photophysical metrics associated with Yale_676sb_ result from the simultaneous introduction of *both* heterocyclic rings *as well as* electron withdrawing dialkyl amino groups (DAGs) into the silicon rhodamine core. When either of these structural features is introduced in isolation, at least one of the three critical photophysical metrics required for two-color SMLM becomes non-optimal. Silicon-rhodamine dyes with only heterocycle-containing dialkyl amino groups (such as HMSiR_indol_, HMSiR_THQ_ and HMSiR_julol_) display long wavelength emission (689 – 716 nm) but resist spirocyclization. As a result, their p*K*_cycl_ values (6.4 – 9.0) are too high to ensure adequate distribution of single-molecule emitters (Figure 2). By contrast, silicon rhodamine dyes with only electron-withdrawing substituents, such as HMSiR_2-FlEt_, display a high quantum yield, but their spirocyclization equilibrium is too favorable, and their p*K*_cycl_ values are too low (Figure 3). By combining these two substitution patterns in Yale_676sb_, the competing effects on p*K*_cycl_ are balanced, while the redshift from the heterocycle-containing DAG is maintained (Figure 3). Moreover, because both the rotational restriction from the heterocycle-containing DAGs and the electron-withdrawing capacity of the 2-fluoroethyl substituent inhibit twisted intramolecular charge transfer (TICT), the quantum yield increase from the latter is not only maintained, but enhanced (0.51 vs 0.59).

As expected, switching from a 2-fluoroethyl to a more electron-withdrawing 2’2-difluoroethyl substituent at the nitrogen in Cal664sb further increases the quantum yield, though at the expense of both p*K*_cycl_ and emission wavelength (Figure 3). This pattern would likely continue with increasingly electron-withdrawing substituents. Despite these blue-shifts, Cal664sb displays a comparable quantum yield to a previously reported and exceptionally bright Si-rhodamine fluorophore (compound **9** in reference 41), but with a >30 nm longer emission maximum.^41^

Finally, we note that while the quantum yield increase relative to HMSiR observed with HMSiR_2-FlEt_ is not as dramatic as that observed with Yale_676sb_, it is comparable to that observed from more commonly used azetidinyl substituents.^16,35,38,46,47^ Being that the former requires only one position at each aniline nitrogen to be substituted, whereas the latter requires two, use of the 2-fluoroethyl substituent may serve as an alternative method for increasing quantum yield of rhodamine derivatives, especially those with more complex DAGs. This approach and others described herein may serve as versatile methods for the preparation of even more greatly enhanced fluorescent labels.

## Acknowledgements

This work was supported by the NIH (1R0GM131372 and 1R35GM134963 to A.S., R01GM118486 to J.B. and D.T., P30DK045735 to the Yale Diabetes Research Center and the Wellcome Trust (grant no. 203285/B/16/Z).

