## Supplementary Figure 1 for "Extremely bright, near-IR emitting spontaneously blinking fluorophores enable ratiometric multicolor nanoscopy in live cells" (from another version of this preprint: version 1, 2021-06-02)

**Table of Contents**


I. **Chemical Synthesis**
   A. **Materials and general information**
   B. **Note on the preparation of NMR samples of $\mathrm{Yale}_{676sb}$ and HMSiR derivatives**
   C. **Supplementary Scheme S1.** Synthetic route to HMSiR
   D. **Supplementary Scheme S2.** Synthetic route to $\mathrm{HMSiR}_{indol}$
   E. **Supplementary Scheme S3.** Synthetic route to $\mathrm{HMSiR}_{julol}$
   F. **Supplementary Scheme S4.** Synthetic route to $\mathrm{HMSiR}_{THQ}$
   G. **Supplementary Scheme S5.** Synthetic route to $\mathrm{Yale}_{676sb}$
   H. **Supplementary Scheme S6.** Synthetic route to $\mathrm{HMSiR}_{2\text{-}FlEt}$
   I. **Supplementary Scheme S7.** Synthetic route to $\mathrm{Cal}_{664sb}$
   J. **Supplementary Scheme S8.** Synthesis of $\mathrm{Yale}_{676sb}$-CA

II. **Biophysical experiments**
   A. **Absorption and fluorescence spectra**
   B. **Plot of Quantum Yield *vs.* Absorption Maxima for Rhodamine-derived Fluorophores (Supplementary Figure 1)**
   C. **Table of Quantum Yield and Absorption Maxima for Rhodamine-derived Fluorophores (Supplementary Table 1)**
   D. **Determination of $pK_{cycl}$ (Supplementary Figure 2)**
   E. **Determination of Quantum Yield (Supplementary Figure 3)**
   F. **Single-molecule kinetics measurements (Supplementary Figure 4)**

III. **Microscopy**
   A. **Cross-talk measurements**
   B. **Labeling Halo-and-SNAP-fusion proteins**
   C. **Super-resolution microscopy and image analysis**

IV. **Supplementary References**

### I. Chemical Synthesis

**A. Materials and general information.** Chemicals used for synthesis were purchased from commercial sources and used without further purification. Flash chromatography was performed using a Teledyne Isco CombiFlash Rf using pre-packed columns with RediSep Rf silica (40–60 µm). $^{1}$H NMR and $^{13}$C NMR spectra were recorded on either an Agilent DD2 NMR spectrometer (400 or 500 MHz for $^{1}$H and 100 or 125 MHz for $^{13}$C) or Bruker NEO500 NMR spectrometer (500 MHz for $^{1}$H and 125 MHz for $^{13}$C). The values of chemical shifts (δ) are reported in p.p.m. relative to the solvent residual signals of $CD_3OD$ (3.31 p.p.m. for $^{1}$H, 49.00 p.p.m. for $^{13}$C), $CDCl_3$ (7.26 p.p.m. for $^{1}$H, 77.36 p.p.m. for $^{13}$C) or $(CD_3)_2O$ (2.05 p.p.m. for $^{1}$H, 29.84, 206.26 p.p.m. for $^{13}$C). Coupling constants (*J*) are reported in Hz. High-resolution mass spectra (HRMS) were recorded on an Agilent 6530QTOF LCMS. HPLC purifications were performed on a reverse-phase column (GL Sciences, Inertsil ODS-3 10 mm × 250 mm and Inertsil ODS-3 20 mm × 250 mm) using a HPLC system composed of a pump (Jasco PU-2080 or PU-2087) and a detector (Jasco MD-2010 or MD-2018).

**B. Note on the preparation of NMR samples of $Yale_{676sb}$ and HMSiR Derivatives.** To simplify the $^{1}$H and $^{13}$C NMR spectra of fluorophores **7**, **11**, **15**, **19**, a single drop of NaOD was added to the NMR sample to fully shift the $pK_{cycl}$ equilibrium of each into the non-emissive closed/OFF form.

**C. Supplementary Scheme S1. Synthesis of HMSiR**

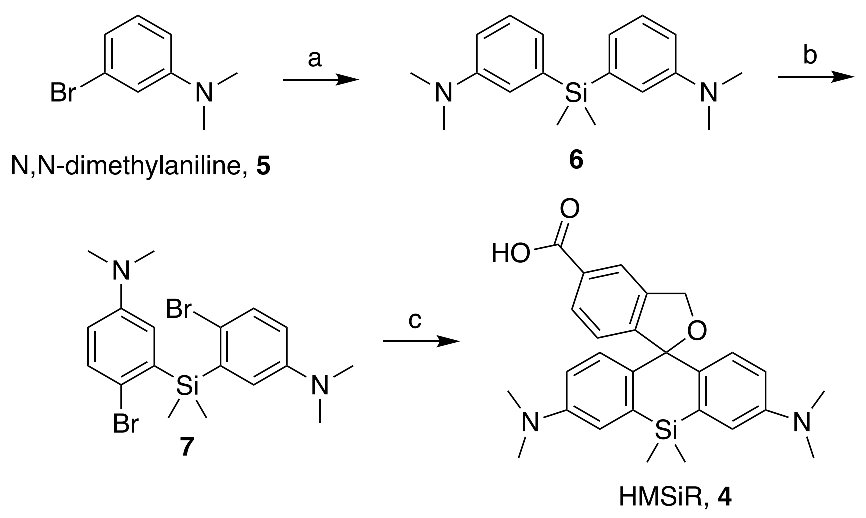


a) n-BuLi, $Me_2SiCl_2$, -78 ˚C to RT, b) NBS, DMF, RT,
c) i. tert-BuLi, **3**, -78 ˚C to RT, ii. TFA, RT

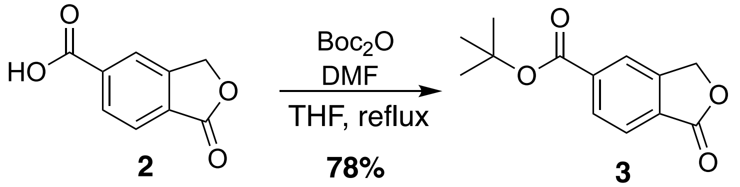


***tert*-butyl-1-oxo-1,3-dihydroisobenzofuran-5-carboxylate 3.** 5’-carboxyphthalide **2** (1.0 g, 2.81 mmol, 1 equiv), $(Boc)_2O$ (3.2 g, 7.02 mmol, 2.5 equiv) and DMAP (140 mg, 0.56 mmol, 0.2 equiv) were dissolved in dry THF (15 mL) and heated to reflux overnight. After cooling to room temperature and evaporation of solvent, the residue was dissolved in $Et_2O$, washed with saturated aqueous $NaHCO_3$, water, and brine, dried over $Na_2SO_4$, filtered, and evaporated. The residue was purified by flash column chromatography on silica (100% hexanes to EtOAc: hexanes 70:30) to obtain **3** as a white powder (78%). $^1$H-NMR (MeOD, 500 MHz) δ 8.21 (s, 1 H), 8.17 (d, *J* = 8.0 Hz, 1 H), 7.96 (d, *J* = 8.0 Hz, 1 H), 5.45 (s, 2 H), 1.64 (s, 9 H). $^{13}$C-NMR (MeOD, 125 MHz) δ 164.5, 147.4, 137.2, 129.6, 128.7, 124.8, 123.3, 82.0, 70.1, 26.9. HRMS calc’d for $C_{13}H_{15}O_4$ $[M+H]^+$ *m/z* 235.0942, found 235.0971.

**3,3'-(dimethylsilanediyl)bis(*N,N*-dimethylaniline) 6.** 3-bromo-N,N-dimethylaniline **5** (5.0 g, 25 mmol, 2.4 equiv) was dissolved in THF (45 mL) in a flame dried round bottom flask and cooled

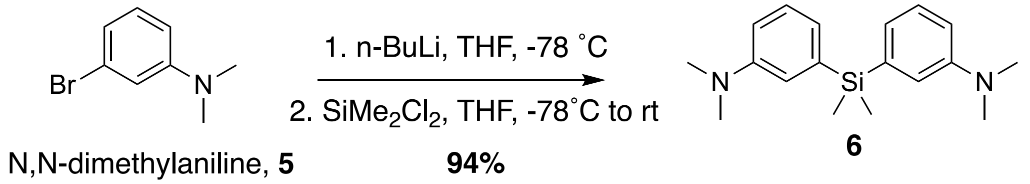


to -78 °C under nitrogen. *n*-Butyllithium (2.5 M in cyclohexane, 10.0 mL, 25 mmol, 2.4 eq) was added and the reaction was stirred at -78 °C for 30 min. Dichlorodimethylsilane (1.26 mL, 10.42 mmol) in THF (5 mL) was then added, the dry ice bath was removed, and the reaction was stirred at room temperature for 3 h. Once no remaining starting material was detectable by TLC analysis, the reaction was quenched with saturated $NH_4Cl$, diluted with water, and extracted twice with EtOAc. The combined organic extracts were washed with brine, dried over anhydrous $Na_2SO_4$, filtered, and concentrated under vacuum. The crude product was purified by flash chromatography on silica (100% hexanes to EtOAc:hexanes 15:85) to yield **6** as a faintly colored oil (94%). $^1$H-NMR ($CDCl_3$, 400 MHz) δ 7.23 (t, *J* = 7.2 Hz, 2 H), 6.92 (m, 4 H), 6.76 (dd, 8.4, 1.6 Hz), 4.60 (t, *J* = 6.0 Hz, 2 H), 4.48 (t, *J* = 6.0 Hz, 2 H), 3.57 (t, *J* = 5.2 Hz, 2 H), 3.51 (t, 5.2 Hz, 2 H), 3.35 (t, *J* = 5.2 Hz, 4 H), 2.91 (s, 12 H), 0.52 (s, 6 H); $^{13}$C NMR ($CDCl_3$, 100 MHz) δ 152.2, 141.2, 130.7, 125.1, 120.6, 115.9, 55.6, 43.0, 0.0. HRMS calc'd for $C_{18}H_{27}N_2Si$ $[M+H]^+$ *m/z* 299.1944, found 299.2323.

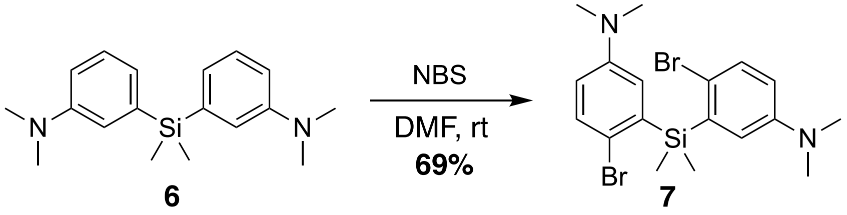


**3,3'-(dimethylsilanediyl)bis(4-bromo-*N*,*N*-dimethylaniline) 7.** Silane **6** (1.0 g, 3.35 mmol) was taken up in DMF (22 mL). N-Bromosuccinimide (1.2 g, 6.70 mmol, 2.0 equiv) was added portionwise over 5 min. The reaction was stirred at room temperature overnight. Product formation was confirmed by LCMS, and then the resulting residue was concentrated in vacuo, diluted with water and extracted twice with EtOAc. The combined organic extracts were washed with water and brine, dried over anhydrous $MgSO_4$, filtered, and concentrated in vacuo. Silica gel chromatography (100% hexanes to EtOAc:hexanes 15:85) afforded **7** as a white solid (69%). $^1$H-NMR ($CDCl_3$, 500 MHz) δ 7.27 (d, *J* = 8.6 Hz, 2 H), 6.77 (d, *J* = 3.3 Hz, 2 H), 6.54 (dd, J = 8.6, 3.2 Hz, 2 H), 2.80 (s, 12 H), 0.69 (s, 6 H). $^{13}$C NMR ($CDCl_3$, 125 MHz) δ 149.7, 139.7, 134.0, 122.8, 117.9, 116.4, 41.6, 0.0. HRMS calc'd for $C_{18}H_{25}Br_2N_2Si$ $[M+H]^+$ *m/z* 455.0154, found 455.0198.

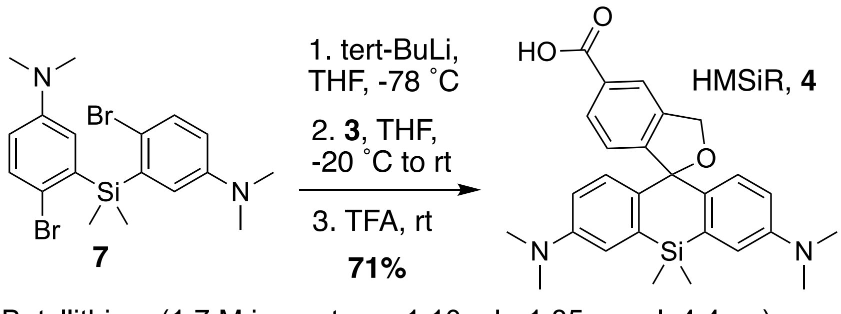


**HMSiR 4.** A solution of dibromide **7** (192 mg, 0.42 mmol) in THF (10 mL) in a flame-dried round bottom flask was cooled to -78 °C under nitrogen. *tert*-Butyllithium (1.7 M in pentane, 1.10 mL, 1.85 mmol, 4.4 eq) was added, and the reaction was stirred at -78 °C for 30 min. The reaction was warmed to -20 °C and then a solution of **3** (216

mg, 0.93 mmol, 2.2 eq) in THF (10 mL) was added dropwise over 30 min. The reaction was allowed to warm to room temperature overnight. Product formation was confirmed by LCMS, and then the reaction was diluted with saturated $NH_4Cl$ and water and extracted with EtOAc (2×). The combined organic extracts were washed with brine, dried over anhydrous $MgSO_4$, filtered, and concentrated *in vacuo*. The resulting residue was redissolved in MeOH (10 mL), and AcOH (100 µL) was added. After stirring for 10 min, the solution was concentrated to dryness and purified by flash chromatography on silica (100% $CH_2Cl_2$ to $CH_2Cl_2$:MeOH 90:10). The resultant material was stirred in TFA for 30 min, concentrated *in vacuo* and re-purified by column chromatography (100% $CH_2Cl_2$ to $CH_2Cl_2$:MeOH 90:10) to afford **HMSiR** as a dark blue solid (71%). $^1$H-NMR (MeOD, 400 MHz) δ 7.90 (s, 1 H), 7.77 (d, $J$ = 8.06 Hz, 1 H), 6.98 (d, $J$ = 8.9 Hz, 2 H), 6.95 (d, $J$ = 2.9 Hz, 2 H), 6.79 (d, $J$ = 8.0 Hz, 1 H), 6.67 (dd, J = 8.9, 2.9 Hz, 2 H), 5.53 (s, 2 H), 2.89 (s, 12 H), 0.51 (d, $J$ = 40.0 Hz, 6 H). $^{13}$C NMR ($CDCl_3$, 125 MHz) δ 176.4, 171.8, 151.9, 151.6, 140.9, 140.1, 139.6, 136.7, 131.0, 125.1, 124.4, 118.7, 116.8, 74.9, 42.1, 1.4, 0.8. HRMS calc'd for $C_{27}H_{31}N_2O_3Si$ $[M]^+$ *m/z* 459.2104, found 459.2214.

### D. Supplementary Scheme S2. Synthesis of HMSiR$_{indol}$

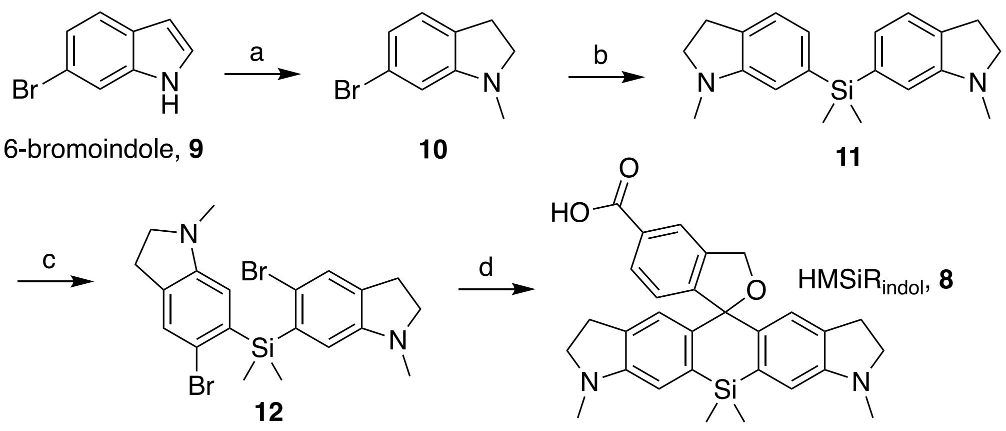


a) formadelhyde, $NaBH_3CN$, r.t., b) n-BuLi, $Me_2SiCl_2$, -78 ˚C to rt, c) NBS, DMF, rt, d) i. tert-BuLi, **3**, -78 ˚C to rt, ii. TFA, rt

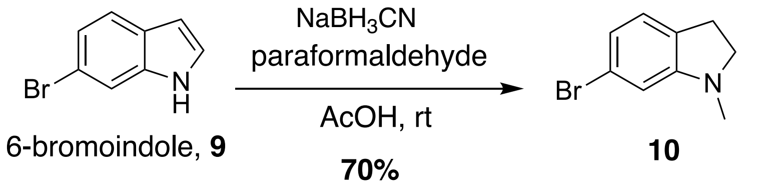


**6-bromo-1-methylindoline 10.** A suspension of 6-bromoindole **9** (2.0 g, 10.20 mmol), paraformaldehyde (3.1 g, 102.02 mmol), and $NaBH_3CN$ (3.2 g, 51.01 mmol) in AcOH (50 mL) was stirred for 3 h at r.t., and the reaction was quenched by adding NaOH. The whole was extracted with $CH_2Cl_2$, and washed with brine. The organic layer was dried over $Na_2SO_4$ and evaporated. Purification of the residue by flash chromatography on silica (100% hexanes to EtOAc:hexanes 15:85) provided **10** as a golden oil (70%). $^1$H NMR ($CDCl_3$, 500 MHz) δ 6.92 (d, *J* = 7.5 Hz, 1 H), 6.79 (dd, *J* = 7.5, 1.7 Hz, 1 H), 6.60 (d, *J* = 1.1 Hz, 1 H), 3.37 (t, *J* = 8.1 Hz, 2 H), 2.92 (t, *J* = 8.1 Hz, 2 H), 2.77 Hz (s, 3 H) . $^{13}$C NMR ($CDCl_3$, 125 MHz) δ 129.5, 125.3, 121.1, 120.6, 110.5, 101.2, 56.2, 35.9, 32.9, 28.2. HRMS calc'd for $C_9H_{11}BrN$ $[M+H]^+$ *m/z* 212.0075, found 212.0149.

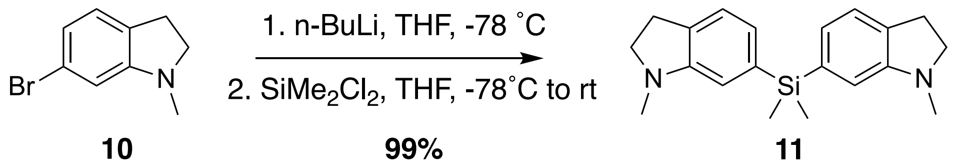


**dimethylbis(1-methylindolin-6-yl)silane 11.** Bromide **10** (1.5 g, 7.07 mmol, 2.4 equiv) was dissolved in THF (13.5 mL) in a flame-dried round bottom flask and cooled to -78 °C under nitrogen. *n*-Butyllithium (2.5 M in cyclohexane, 2.83 mL, 7.07 mmol, 2.4 eq) was added, and the reaction was stirred at -78 °C for 30 min. Dichlorodimethylsilane (0.356 mL, 2.95 mmol) in THF (5 mL) was then added, the dry ice bath was removed, and the reaction was stirred at room temperature for 3 h. It was subsequently quenched with saturated $NH_4Cl$, diluted with water, and extracted with EtOAc (2×). The combined organic phase was washed with brine, dried over anhydrous Na2SO4, filtered, and concentrated under vacuum. The crude product was purified by flash chromatography on silica (100% hexanes to EtOAc:hexanes 15:85) to yield **11** as a colorless oil (99%). $^1$H-NMR ($CDCl_3$, 400 MHz) δ 7.07 (d, J = 7.1 Hz, 2 H), 6.85 (d, J = 7.3 Hz, 2 H), 6.64 (s, 2 H), 3.27 (t, 7.90 Hz, 4 H), 2.92 (t, 8.20 Hz, 4 H), 2.73 (s, 6 H), 0.49 (s, 6H). HRMS calc'd for $C_{20}H_{27}N_2Si$ $[M+H]^+$ *m/z* 323.1944, found 323.1925.

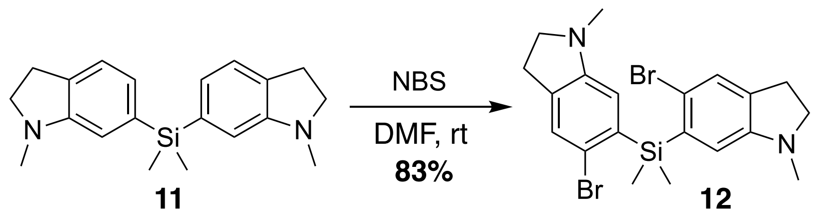


**bis(5-bromo-1-methylindolin-6-yl)dimethylsilane 12.** Silane **11** (1.05 g, 3.26 mmol) was taken up in DMF (10 mL). N-Bromosuccinimide (1.16 g, 6.52 mmol, 2 eq) was dissolved in 10 mL DMF and added dropwise over 5 min at 0 ˚C. The reaction was then stirred at room temperature overnight. Product formation was confirmed by LCMS, and then the resulting residue was concentrated in vacuo, diluted with water and extracted twice with EtOAc. The combined organic extracts were washed with water and brine, dried over anhydrous $MgSO_4$, filtered, and concentrated in vacuo. Silica gel chromatography (100% hexanes to EtOAc:hexanes 15:85) afforded **12** as a reddish brown solid (83%). $^1$H NMR ($CDCl_3$, 400 MHz) δ 7.18 (s, 2 H), 6.54 (s, 2 H), 3.27 (t, J = 8.1 Hz, 4 H), 2.91 (t, J = 8.2 Hz, 4

H), 2.68 (s, 6 H), 0.70 (s, 6 H). HRMS calc'd for $C_{20}H_{25}Br_2N_2Si$ $[M+H]^+$ *m/z* 479.0154, found 479.0185. $^{13}C$ NMR ($CDCl_3$, 125 MHz) δ 137.7, 134.9, 129.5, 118.9, 116.1, 100.9, 56.8, 37.1, 29.1, 0.0. HRMS calc'd for $C_{20}H_{25}Br_2N_2Si$ $[M+H]^+$ *m/z* 479.0154, found 479.0185.

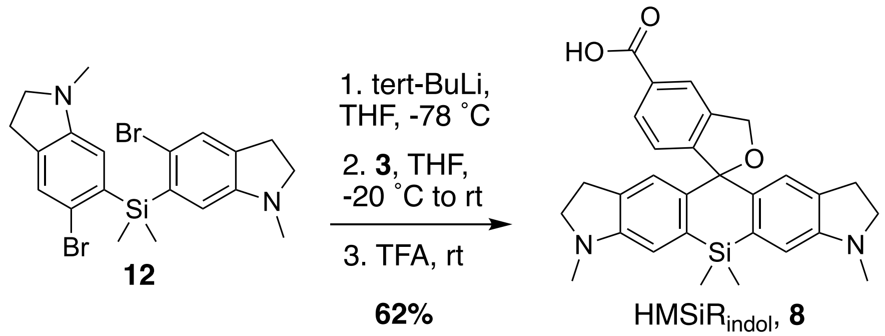


**HMSiR$_{indol}$ 8.** A solution of dibromide **8** (428 mg, 0.89 mmol) in THF (16 mL) was cooled to -78 °C under nitrogen. *tert*-Butyllithium (1.7 M in pentane, 2.29 mL, 3.9 mmol, 4.4 eq) was added, and the reaction was stirred at -78 °C for 30 min. A solution of **3** (435 mg, 1.78 mmol, 2.2 eq) in THF (16 mL) was added dropwise over 30 min. The reaction was allowed to warm to room temperature overnight. Product formation was confirmed by LCMS, and it was subsequently diluted with saturated NH4Cl and water and extracted with EtOAc (2×). The combined organic extracts were washed with brine, dried over anhydrous $MgSO_4$, filtered, and concentrated *in vacuo*. The resulting residue was redissolved in MeOH (10 mL), and AcOH (100 µL) was added. After stirring for 10 min, the solution was concentrated to dryness and purified by flash chromatography on silica (100% $CH_2Cl_2$ to $CH_2Cl_2$:MeOH 90:10). The resultant material was stirred in TFA for 30 min, concentrated *in vacuo* and re-purified by column chromatography (100% $CH_2Cl_2$ to $CH_2Cl_2$:MeOH 90:10) to afford **HMSiR$_{indol}$** as a bluish green solid (62%). $^1H$-NMR (MeOD, 500 MHz) δ 8.39 (s, 1 H), 8.06 (dd, 7.6, 1.8 Hz, 1 H), 7.20 (d, *J* = 7.8 Hz, 1 H), 7.13 (s, 2 H), 6.59 (s, 2 H), 4.34 (s, 2 H), 3.80 (t, *J* = 8.1 Hz, 4H), 3.21 (s, 6 H), 2.89 (t, *J* = 8.1 Hz, 4H), 0.55 (d, *J* = 5.7 Hz, 6 H). $^{13}C$ NMR ($CDCl_3$, 125 MHz) δ 167.8, 162.6, 157.0, 150.2, 142.5, 140.0, 134.9, 133.3, 131.9, 131.1, 129.1, 127.9, 127.7, 114.6, 60.4, 54.2, 32.3, 25.7, -2.6, -2.9. HRMS calc'd for $C_{29}H_{31}N_2O_3Si$ $[M]^+$ *m/z* 483.2104, found 483.1947.

E. **Supplementary Scheme S3.** Synthesis of $HMSiR_{julol}$.

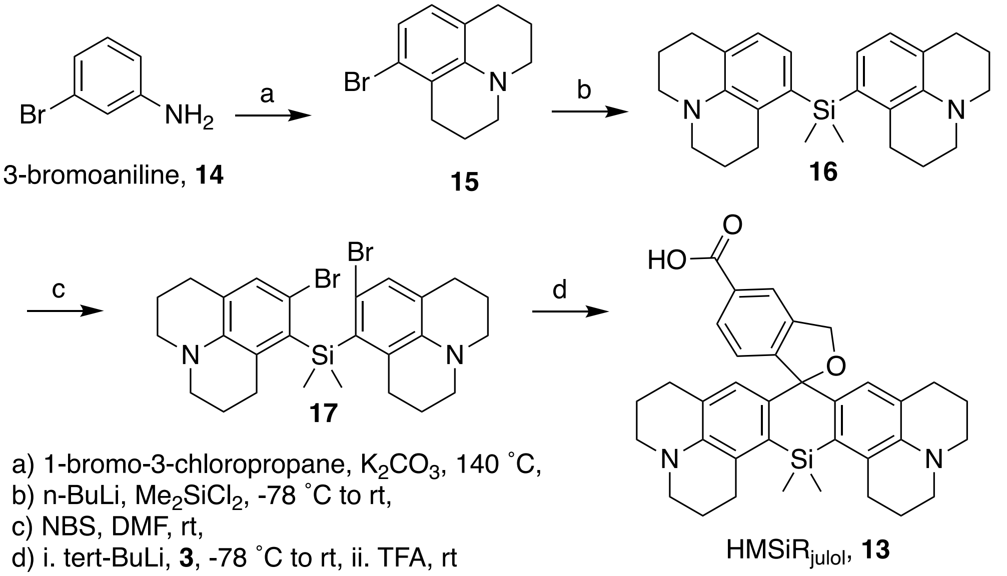


a) 1-bromo-3-chloropropane, $K_2CO_3$, 140 ˚C,
b) n-BuLi, $Me_2SiCl_2$, -78 ˚C to rt,
c) NBS, DMF, rt,
d) i. tert-BuLi, **3**, -78 ˚C to rt, ii. TFA, rt

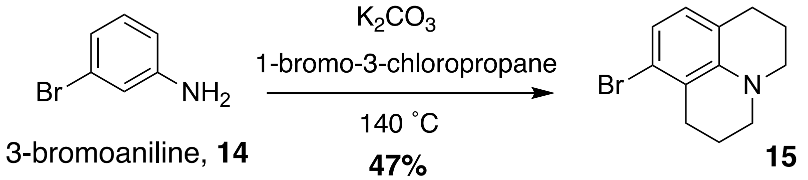


**8-bromojulolidine 15.** A round-bottom flask was charged with 3-bromoaniline **14** (3.95 g, 22.96 mmol), 1-bromo-3-chloropropane (16.27 mL, 103.33 mmol, 4.5 eq.), and $K_2CO_3$ (6.35 g, 45.93 mmol, 2.0 eq.). After stirring at 140 °C for 48 h, the reaction was cooled to room temperature, transferred to a separatory funnel, and extracted with $CH_2Cl_2$ 3x. The combined organic fractions were dried ($Na_2SO_4$), filtered, and concentrated under reduced pressure. Residual 1-bromo-3-chloropropane was removed via flash column chromatography (100% hexanes to EtOAc:hexanes 5:95). The crude mixture (containing some product, and some dialkylated intermediate) was dissolved in DMF (5.0 mL) and stirred at 160 °C for 48 hrs. After cooling to room temperature, the solution was concentrated under reduced pressure, washed with water, and extracted with $CH_2Cl_2$ 3x. The combined organic fractions were dried over $Na_2SO_4$, filtered, and concentrated under reduced pressure. The crude residue was purified via flash chromatography on silica (100% hexanes to EtOAc:hexanes 5:95 very slowly) to afford **15** as a reddish oil (47%). $^1H$ NMR (500 MHz, $CDCl_3$) δ 6.70 (d, J = 8.0 Hz, 1H), 6.57 (d, J = 8.0 Hz, 1H), 3.05 (m, 4H), 2.70 (t, J = 6.7 Hz, 2H), 2.63 (t, J = 6.5 Hz, 2H), 1.9 (m, 4H). $^{13}C$ NMR (125 MHz, CDCl3) δ 144.5, 127.8, 122.9, 120.7, 120.6, 119.5, 50.0, 49.7, 28.8, 27.6, 21.9, 21.8. HRMS calc'd for $C_{12}H_{15}BrN$ $[M+H]^+$ *m/z* 252.0388, found 252.0566.

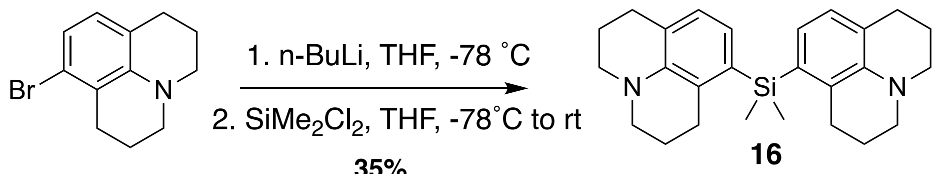

15



16

**dimethylbis(2,3,6,7-tetrahydro-1*H*,5*H*-pyrido[3,2,1-*ij*]quinolin-8-yl)silane 16.** Bromide 15 (1.513 g, 6.0 mmol, 2.4 equiv) was dissolved in THF (12 mL) in a flame dried round bottom flask and cooled to -78 °C under nitrogen. *n*-Butyllithium (2.5 M in cyclohexane, 2.4 mL, 6.0 mmol, 2.4 eq) was added, and the reaction was stirred at -78 °C for 30 min. Dichlorodimethylsilane (0.304 mL, 2.5 mmol) in THF (1.2 mL) was then added dropwise, the dry ice bath was removed, and the reaction was slowly warmed up to room temperature for 3 h. It was subsequently quenched with saturated $NH_4Cl$, diluted with water, and extracted with EtOAc (2×). The combined organic phase was washed with brine, dried over anhydrous $Na_2SO_4$, filtered, and concentrated under vacuum. The crude product was purified by flash chromatography on silica (100% hexanes to EtOAc:hexanes 5:95) to yield 16 as a colorless oil (352 mg, 35%).$^1$H-NMR ($CDCl_3$, 400 MHz) δ 6.77 (d, *J* = 7.4 Hz, 2H), 6.69 (d, *J* = 7.4 Hz, 2H), 3.13 (t, *J* = 5.5 Hz, 4H), 3.07 (t, *J* = 5.8 Hz, 4H), 2.76 (t, *J* = 6.5 Hz, 4H), 2.67 (t, *J* = 6.4 Hz, 4H), 1.96 (dt, *J* = 12.2, 6.4 Hz, 4H), 1.85 (dt, *J* = 12.4, 6.3 Hz, 4H), 0.51 (s, 6H). $^{13}$C NMR ($CDCl_3$, 125 MHz) δ142.9, 135.3, 127.7, 126.7, 123.3, 122.9, 50.7, 50.2, 28.7, 28.2, 22.5, 22.2, 0.2. HRMS calc'd for $C_{26}H_{35}N_2Si$ $[M+H]^+$ *m/z* 403.2564, found: 403.2519.

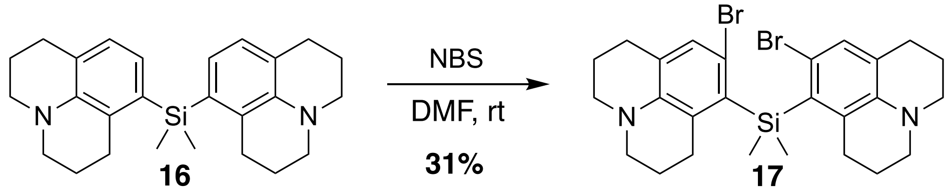

16



17

**bis(9-bromo-2,3,6,7-tetrahydro-1H,5H-pyrido[3,2,1-ij]quinolin-8-yl)dimethylsilane 17.** Silane 16 (350.2 mg, 0.87 mmol) was taken up in DMF (4 mL). N-Bromosuccinimide (309.6 mg, 1.74 mmol, 2 eq) was dissolved in 2 mL DMF and added dropwise over 5 min at 0 ˚C. The reaction was then stirred at room temperature overnight. Product formation was confirmed by LCMS, and then the resulting residue was concentrated in vacuo, diluted with water and extracted twice with EtOAc. The combined organic extracts were washed with brine(3x), dried over anhydrous $Na_2SO_4$, filtered and concentrated in vacuo. Silica gel chromatography (100% hexanes to EtOAc:hexanes 5:95) afforded 17 as a white solid (31%). $^1$H NMR ($CDCl_3$, 500 MHz) δ 6.94 (s, 2H), 3.10 (t, *J* = 5.6 Hz, 4H), 3.04 (t, *J* = 6.0 Hz, 4H), 2.82 – 2.57 (m, 8H), 1.91 (dt, *J* = 12.3, 6.2 Hz, 4H), 1.81 (dt, *J* = 12.4, 6.1 Hz, 4H), 0.78 (s,

6H). $^{13}C$ NMR ($CDCl_3$, 125 MHz) δ 142.0, 137.7, 131.9, 129.5, 124.7, 115.5, 50.6, 49.7, 29.2, 27.8, 22.5, 21.9, 7.7. HRMS calc'd for $C_{26}H_{33}Br_2N_2Si$ $[M+H]^+$ *m/z* 559.0774, found: 559.0749.

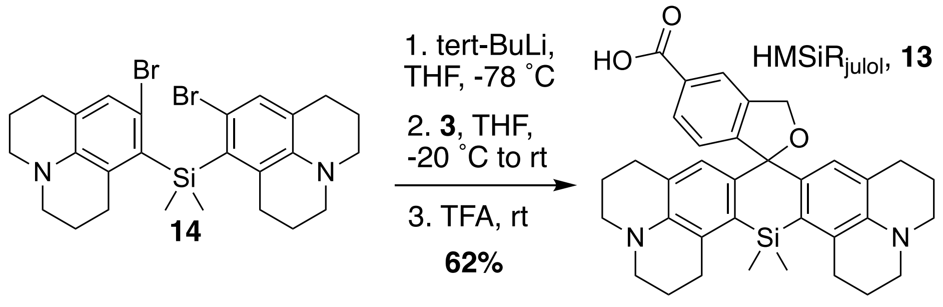


**HMSiR$_{julol}$ 13.** A solution of dibromide **14** (56 mg, 0.10 mmol) in THF (2 mL) was cooled to -78 °C under nitrogen. *tert*-Butyllithium (1.7 M in pentane, 0.26 mL, 0.44 mmol, 4.4 eq) was added, and the reaction was stirred at -78 °C for 30 min. A solution of **3** (51.5 mg, 0.22 mmol, 2.2 eq) in THF (2.2 mL) was added dropwise over 30 min. The reaction was allowed to warm to room temperature overnight. Product formation was confirmed by LCMS, and the reaction mixture was subsequently quenched with saturated $NH_4Cl$ and extracted with EtOAc (2×). The combined organic extracts were washed with brine, dried over anhydrous $Na_2SO_4$, filtered, and concentrated *in vacuo*. The resulting residue was redissolved in MeOH (2 mL), and TFA (50 µL) was added. After stirring for 10 min, the solution was concentrated to dryness. The resultant material was subsequently stirred in TFA (1 mL) for 30 min, concentrated *in vacuo* and purified by column chromatography (100% $CH_2Cl_2$ to $CH_2Cl_2$:MeOH 95:5) to afford HMSiR$_{julol}$ as a bluish green solid (42.1 mg, 62%). $^1H$-NMR (MeOD with 0.1% NaOD, 500 MHz) δ 7.82 (d, *J* = 1.4 Hz, 1H), 7.63 (dd, *J* = 8.0, 1.5 Hz, 1H), 6.70 (s, 2H), 6.58 (d, *J* = 8.0 Hz, 1H), 5.51 (s, 2H), 3.16 (t, *J* = 6.0 Hz, 4H), 3.11 (td, *J* = 5.0, 2.8 Hz, 4H), 2.95 – 2.82 (m, 4H), 2.66 – 2.44 (m, 4H), 1.99 (tt, *J* = 6.7, 4.7 Hz, 4H), 1.86 (tt, *J* = 6.8, 4.7 Hz, 4H), 0.73 (s, 3H), 0.55 (s, 3H). $^{13}C$ NMR ($CDCl_3$, 125 MHz) δ 175.4, 163.2, 153.4, 143.0, 139.3, 138.0, 136.4, 130.4, 129.9, 127.7, 127.0, 125.3, 123.3, 123.0, 95.8, 75.0, 51.5, 51.0, 30.3, 29.4, 23.5, 23.2, 2.5, 2.1. HRMS calc'd for $C_{35}H_{39}N_2O_3Si$ $[M]^+$ *m/z* 563.2724, found: 563.2701.

**F. Supplementary Scheme S4.** Synthesis of $HMSiR_{THQ}$.

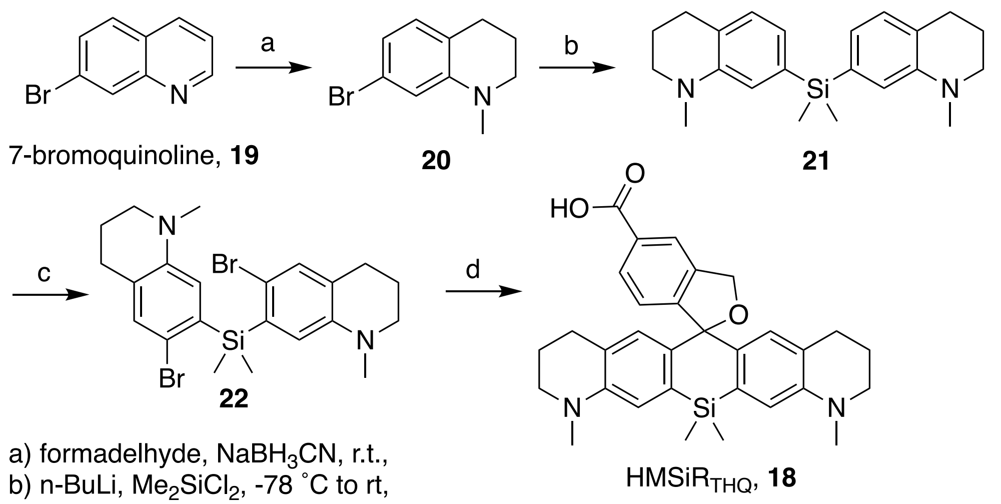


a) formadelhyde, $NaBH_3CN$, r.t.,
b) n-BuLi, $Me_2SiCl_2$, -78 ˚C to rt,
c) NBS, DMF, rt,
d) i. tert-BuLi, **3**, -78 ˚C to rt, ii. TFA, rt

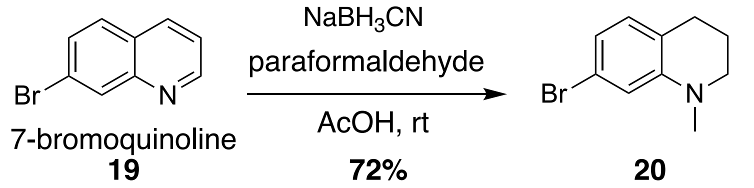


**7-bromo-1-methyl-1,2,3,4-tetrahydroquinoline 20.** A suspension of 7-bromoquinoline **19** (5.2 g, 25.0 mmol), paraformaldehyde (7.5 g, 250.0 mmol, 10 equiv), and $NaBH_3CN$ (7.9 g, 125.0 mmol, 5.0 equiv) in AcOH (125 mL) was stirred for 3 h at RT, and the reaction was quenched by adding NaOH. The whole was extracted with $CH_2Cl_2$, and washed with brine. The organic layer was dried over $Na_2SO_4$ and evaporated. Purification of the residue by flash chromatography on silica (100% hexanes to EtOAc:hexanes 15:85) provided **20** as a golden oil (70%). $^1$H NMR ($CDCl_3$, 400 MHz) δ 6.81 (d, *J* = 7.8 Hz, 1 H), 6.72 (m, 2 H), 3.25 (t, *J* = 5.7 Hz, 2 H), 2.90 Hz (s, 3 H), 2.71 (t, *J* = 6.4 Hz, 2 H), 1.98 (m, 2 H) . $^{13}$C NMR ($CDCl_3$, 125 MHz) δ 130.6, 129.9, 121.7, 120.6, 118.8, 113.6, 50.9, 39.1, 27.3, 22.0. HRMS calc'd for $C_{10}H_{13}BrN$ $[M+H]^+$ *m/z* 226.0231, found 226.0105.

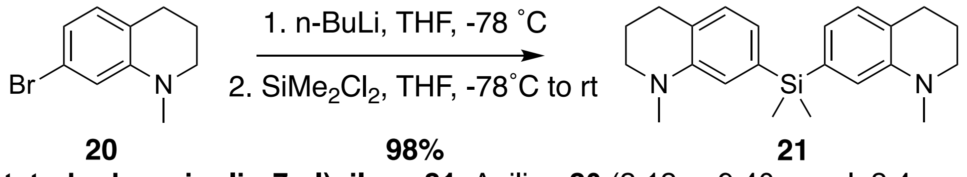


**Dimethylbis(1-methyl-1,2,3,4-tetrahydroquinolin-7-yl)silane 21.** Aniline **20** (2.12 g, 9.40 mmol, 2.4 equiv) was dissolved in THF (18 mL) in a flame-dried round bottom flask and cooled to -78 °C under nitrogen. *n*-Butyllithium (2.5 M in cyclohexane, 3.60 mL, 9.40 mmol, 2.4 eq) was added, and the reaction was stirred at -78 °C for 30 min. Dichlorodimethylsilane (0.50 mL, 3.90 mmol) in THF (5 mL) was then added, the dry ice bath was removed, and the reaction was stirred at room temperature for 3 h. It was subsequently quenched with saturated $NH_4Cl$, diluted with water, and extracted with EtOAc (2×). The combined organic phase was washed with brine, dried over anhydrous $Na_2SO_4$, filtered, and concentrated under vacuum. The crude product was purified by flash chromatography on silica (100% hexanes to EtOAc:hexanes 15:85) to yield **21** as a colorless oil (99%). $^1$H NMR ($CDCl_3$, 400 MHz) δ 6.92 (d, J = 7.2 Hz, 2 H), 6.78 (m, 4 H), 4.02 (t, J = 5.6, 4 H), 2.85 (s, 6 H), 2.74 (t, 6.1 Hz, 4 H), 4.10 (m, 4 H), 0.47 (s, 6 H). $^{13}$C NMR ($CDCl_3$, 125 MHz) δ 148.1, 138.72, 130.4, 125.9, 124.5, 118.6, 53.5, 41.2, 29.9, 24.4, 0.00. HRMS calc'd for $C_{22}H_{31}N_2Si$ $[M+H]^+$ *m/z* 351.2257, found 351.2358.

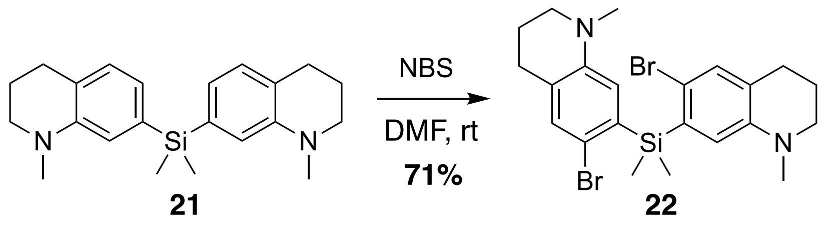


**bis(6-bromo-1-(2,2-difluoroethyl)-1,2,3,4-tetrahydroquinolin-7-yl)dimethylsilane 22.** Silane **21** (89 mg, 0.25 mmol) was taken up in DMF (0.8 mL). N-Bromosuccinimide (90.37 mg, 0.51 mmol, 2.0 eq) was dissolved in 0.8 mL DMF and added dropwise over 5 min at 0 ˚C. The reaction was then stirred at room temperature overnight. Product formation was confirmed by LCMS, and then the resulting residue was concentrated in vacuo, diluted with water and extracted twice with EtOAc. The combined organic extracts were washed with water and brine, dried over anhydrous $MgSO_4$, filtered, and concentrated in vacuo. Silica gel chromatography (100% hexanes to EtOAc:hexanes 15:85) afforded **22** as a reddish brown solid (69%). $^1$H-NMR ($CDCl_3$, 400 MHz) δ 7.07 (s, 2 H), 6.67 (s, 2 H), 3.16 (t, J = 5.6 Hz, 4 H), 2.77 (s, 6 H), 2.70 (t, J = 6.4 Hz, 4 H), 1.93 (p, J = 6.0 Hz, 4 H), 0.69 (s, 6 H). $^{13}$C NMR ($CDCl_3$, 125 MHz) δ 145.9, 137.1, 133.3, 127.0, 120.9, 117.01, 52.0, 39.8, 28.2, 22.9, 0.00. HRMS calc'd for $C_{22}H_{29}Br_2N_2Si$ $[M+H]^+$ *m/z* 507.0467, found 507.0157.

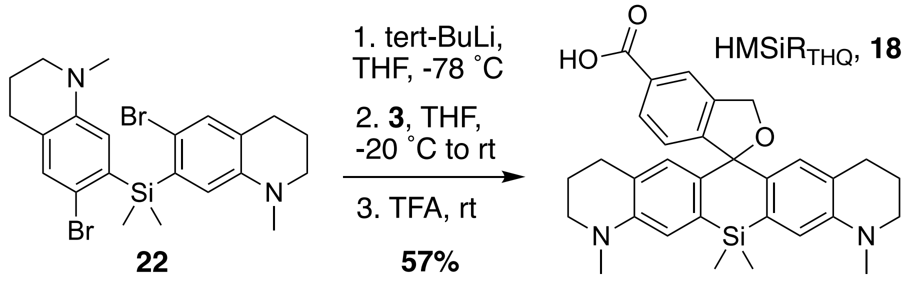


**HMSiR$_{THQ}$ 18.** A solution of dibromide **22** (428 mg, 0.89 mmol) in THF (16 mL) was cooled to -78 °C under nitrogen. *tert*-Butyllithium (1.7 M in pentane, 2.29 mL, 3.9 mmol, 4.4 eq) was added, and the reaction was stirred at -78 °C for 30 min. A solution of **3** (435 mg, 1.78 mmol, 2.2 eq) in THF (16 mL) was added dropwise over 30 min. The reaction was allowed to warm to room temperature overnight. Product formation was confirmed by LCMS, and it was subsequently diluted with saturated $NH_4Cl$ and water and extracted with EtOAc (2×). The combined organic extracts were washed with brine, dried over anhydrous $MgSO_4$, filtered, and concentrated *in vacuo*. The resulting residue was redissolved in MeOH (10 mL), and AcOH (100 µL) was added. After stirring for 10 min, the solution was concentrated to dryness and purified by flash chromatography on silica (100% $CH_2Cl_2$ to $CH_2Cl_2$:MeOH 90:10). The resultant material was stirred in TFA for 30 min, concentrated *in vacuo* and re-purified by column chromatography (100% $CH_2Cl_2$ to $CH_2Cl_2$:MeOH 90:10) to afford **$\mathbf{HMSiR_{THQ}}$** as a bluish green solid (57%). $^1$H-NMR (MeOD, 500 MHz) δ 7.97 (s, 1 H), 7.90 (s, 1 H), 7.75 (d, *J* = 8.0 Hz, 1 H), 6.76 (s, 2 H), 6.69 (s, 2 H), 5.38 (s, 2 H), 3.17 (t, *J* = 5.3 Hz), 2.90 (s, 6 H), 4.57 (m, 4 H), 1.88 (m, 4 H), 0.48 (d, *J* = 48.0 Hz, 6 H). $^{13}$C NMR (151 MHz, MeOD) δ 173.94, 150.07, 145.34, 138.35, 137.06, 131.61, 128.47, 128.15, 124.70, 122.34, 121.81, 114.33, 92.83, 72.63, 69.09, 51.04, 48.16, 48.02, 47.87, 47.73, 47.59, 47.45, 47.31, 47.16, 37.96, 28.12, 27.74, 22.14, -1.18, -1.35. HRMS calc'd for $C_{31}H_{35}N_2O_3Si$ $[M]^+$ *m/z* 511.2417, found 511.2417.

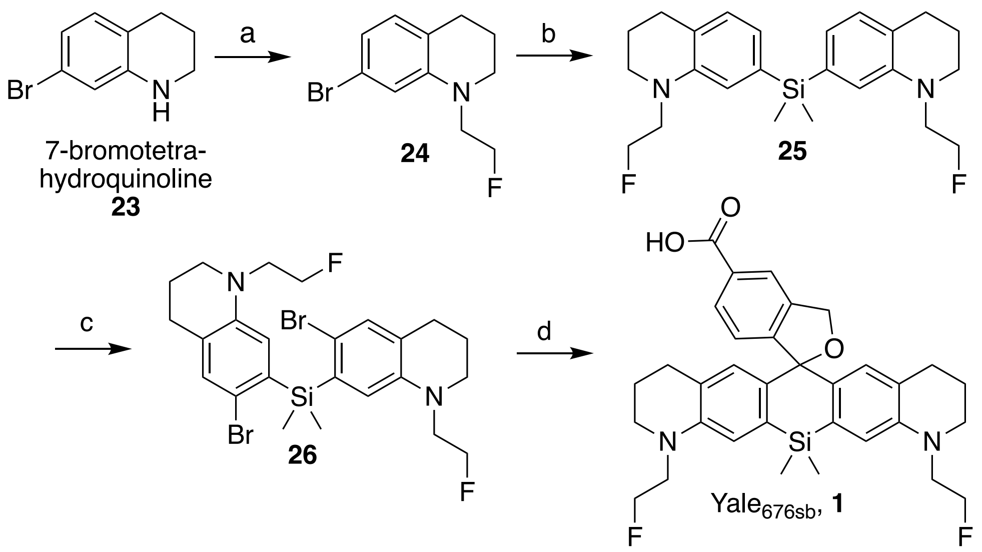


**G. Supplementary Scheme S5.** Synthesis of $Yale_{676sb}$.

a) 1-fluoro-2-iodoethane, $K_2CO_3$, DMF, 70˚C, b) n-BuLi, $Me_2SiCl_2$, -78 ˚C to rt, c) NBS, DMF, rt, d) i. tert-BuLi, **3**, -78 ˚C to rt, ii. TFA, rt

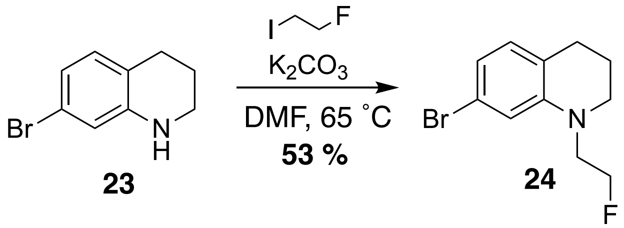


**7-bromo-1-(2-fluoroethyl)-1,2,3,4-tetrahydroquinoline 24.** Potassium carbonate (1.33 g, 9.6 mmol, 1.5 equiv) was added to a solution of 7-bromo-1,2,3,4-tetrahydroquinoline **23** (1.36 g, 6.4 mmol, 1.0 equiv) in DMF (5.5 mL). 1-bromo-2-fluoroethane (7.7 mmol, 1.2 equiv) was added, and the reaction was stirred at 65 °C for 18 h. The mixture was allowed to cool to room temperature, diluted with water, and acidified with 1 M HCl. The aqueous phase was extracted with ethyl acetate, and the combined organic phases were washed 3x with water and brine and dried over $Na_2SO_4$. The solvent was evaporated under vacuum, and the crude product was purified by flash column chromatography on silica (100% hexanes to EtOAc:hexanes 5:95) to yield **24** (53%) as a yellow oil. $^1$H-NMR ($CDCl_3$, 400 MHz) δ 6.78 (d, *J* = 7.7 Hz, 1 H), 6.68 (dd, *J* = 7.9, 1.7 Hz, 1 H), 6.61 (d, *J* = 1.8 Hz, 1 H), 4.66 (t, *J* = 5.2 Hz, 1 H), 4.54 (t, *J* = 5.2 Hz, 1 H), 3.58 (t, *J* = 5.2 H, 1 H), 3.52 (t, *J* = 5.2 H, 1 H), 3.35 (t, *J* = 5.8 Hz, 2 H), 2.68 (t, *J* = 6.4, 2 H), 1.92 (m, *J* = 5.8, 6.2 Hz, 2 H). $^{13}$C NMR ($CDCl_3$, 125 MHz) δ 146.0, 130.5, 121.4, 120.6, 118.7, 112.9, 82.1, 80.7, 51.5, 51.3, 50.3, 27.7, 21.9. HRMS calc'd for $C_{10}H_{13}BrN$ $[M+H]^+$ *m/z* 226.0231, found 226.0105.

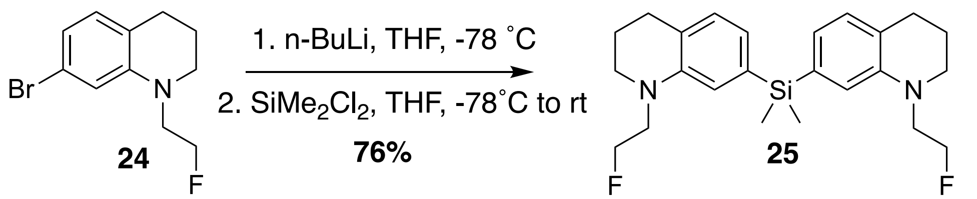


**bis(1-(2-fluoroethyl)-1,2,3,4-tetrahydroquinolin-7-yl)dimethylsilane 25.** Aniline **24** (398 mg, 1.54 mmol, 2.4 equiv) was dissolved in THF (3.6 mL) in a flame-dried round bottom flask and cooled to -78 °C under nitrogen. *n*- Butyllithium (2.5 M in cyclohexane, 0.62 mL, 1.54 mmol, 2.4 eq) was added, and the reaction was stirred at -78 °C for 30 min. Dichlorodimethylsilane (0.08 mL, 0.64 mmol) in THF (0.4 mL) was then added, the dry ice bath was removed, and the reaction was stirred at room temperature for 3 h. It was subsequently quenched with saturated $NH_4Cl$, diluted with water, and extracted with EtOAc (2×). The combined organic phase was washed with brine, dried over anhydrous $Na_2SO4$, filtered, and concentrated under vacuum. The crude product was purified by flash chromatography on silica (100% hexanes to EtOAc:hexanes 15:85) to yield **25** as a bronze oil (76%). $^1$H-NMR ($CDCl_3$, 400 MHz) δ 6.93 (d, *J* = 7.2 Hz, 2 H), 6.75 (d, *J* = 7.3 Hz, 2 H), 6.65 (s, 2 Hz), 4.60 (t, *J* = 6.0 Hz, 2 H), 4.48 (t, *J* = 6.0 Hz, 2 H), 3.57 (t, *J* = 5.2 Hz, 2 H), 3.51 (t, 5.2 Hz, 2 H), 3.35 (t, *J* = 5.2 Hz, 4 H), 2.74 (t, *J* = 6.3 Hz, 4 H), 1.93 (m, 4 H), 0.45 (s, 6 H); $^{13}$C NMR ($CDCl_3$, 125 MHz) δ 146.4, 138.9, 131.1, 126.0, 124.5, 118.3, 84.4, 83.1, 53.9, 53.7, 52.9, 30.27, 24.27, 0.0. HRMS calc'd for $C_{24}H_{33}F_2N_2Si$ $[M+H]^+$ *m/z* 415.2381, found 415.2366.

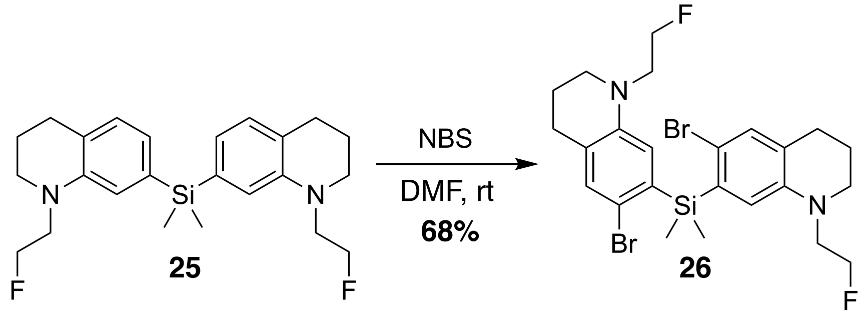

**bis(6-bromo-1-(2-fluoroethyl)-1,2,3,4-tetrahydroquinolin-7-yl)dimethylsilane 26.** Silane **25** (35.5 mg, 0.086 mmol) was taken up in DMF (0.8 mL). N-Bromosuccinimide (30.5 mg, 0.17 mmol, 2 eq) was dissolved in 0.8 mL DMF and added dropwise over 5 min at 0 ˚C. The reaction was then stirred at room temperature overnight. Product formation was confirmed by LCMS, and then the resulting residue was concentrated in vacuo, diluted with water and extracted twice with EtOAc. The combined organic extracts were washed with water and brine, dried over anhydrous $MgSO_4$, filtered, and concentrated in vacuo. Silica gel chromatography (100% hexanes to EtOAc:hexanes 15:85) afforded **26** as a reddish brown solid (68%). $^1$H-NMR ($CDCl_3$, 400 MHz) δ 7.09 (s, 2 H), 6.51 (s, 2 H), 4.53 (t, *J* = 5.3 Hz, 2 H), 4.41 (*J* = 5.3 Hz, 2 H), 3.47 (t, *J* = 5.2 Hz, 2 H), 3.41 (t, *J* = 5.2 Hz, 2H), 3.32 (t, *J* = 5.5 Hz, 4 H), 2.71 (t, *J* = 6.3 Hz, 4 H), 1.91 (m, 4 H), 0.69 (s, 6 H). $^{13}$C NMR ($CDCl_3$, 125 MHz) δ 144.1 (C), 137.4 (CH), 133.7 (CH), 126.9 (C), 120.3 (C), 116.7 (C), 82.9 ($CH_2F$), 81.5 ($CH_2F$), 52.6 ($CH_2$), 52.4 ($CH_2$), 51.4 ($CH_2$), 28.4 ($CH_2$), 22.7 ($CH_2$), 0.00 ($CH_3$). HRMS calc'd for $C_{24}H_{31}Br_2F_2N_2Si$ $[M+H]^+$ *m/z* 571.0591, found 571.0268.

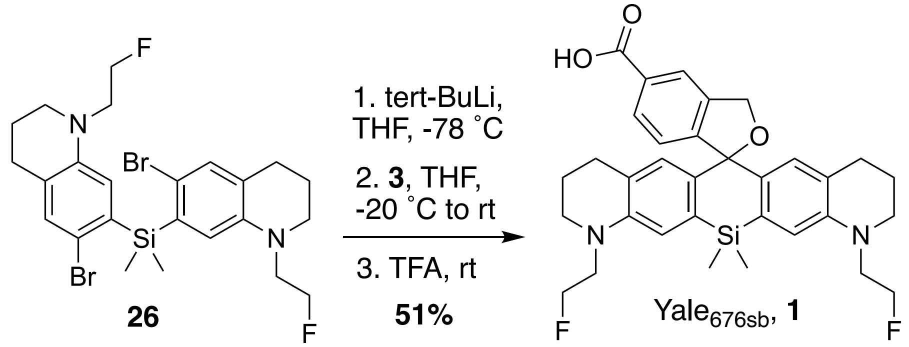

**Yale$_{676sb}$ 1.** A solution of dibromide **26** (52.5 mg, 0.092 mmol) in THF (2.5 mL) was cooled to -78 °C under nitrogen. *tert*-Butyllithium (1.7 M in pentane, 0.24 mL, 0.404 mmol, 4.4 eq) was added, and the reaction was stirred at -78 °C for 30 min. A solution of **3** (47.3 mg, 0.202 mmol, 2.2 eq) in THF (2.5 mL) was added dropwise over 30 min. The reaction was allowed to warm to room temperature overnight. Product formation was confirmed by LCMS, and it was subsequently diluted with saturated $NH_4Cl$ and water and extracted with EtOAc (2×). The combined organic extracts were washed with brine, dried over anhydrous MgSO4, filtered, and concentrated *in vacuo.* The resulting residue was redissolved in MeOH (3 mL), and AcOH (100 μL) was added. After stirring for 10 min, the solution was concentrated to dryness and purified by flash chromatography on silica (100% $CH_2Cl_2$ to $CH_2Cl_2$:MeOH 90:10). The resultant material was stirred in TFA for 30 min, concentrated *in vacuo* and re-purified by column chromatography (100% $CH_2Cl_2$ to $CH_2Cl_2$:MeOH 90:10) to afford **Yale$_{676sb}$** as a bluish green solid (51%). $^1$H-NMR (MeOD, 500 MHz) δ 7.93 (s, 1 H), 7.79 (d, *J* = 7.9 Hz, 1 H),6.76 (d, *J* = 8.0 Hz, 1 H), 6.71 (s, 2 H), 6.65 (s, 2 H), 5.36 (s, 2 H), 4.62 (dt, *J* = 47.9, 4.5 Hz, 4), 3.61 (m, 4 H), 3.34 (m, 4 H), 2.53 (m, 4 H), 1.83 (m, 4 H), 0.43 (d, *J* = 44.8 Hz, 6 H). $^{13}$C NMR ($CDCl_3$, 125 MHz) δ 174.0, 149.9, 143.6, 137.8, 137.2, 137.0, 131.7, 128.6, 128.5, 124.2, 122.5, 121.8, 114.1, 92.8, 82.1, 80.8, 72.5, 51.2, 51.1, 50.2, 28.1, 22.0, -1.2, -1.5. HRMS calc'd for $C_{33}H_{37}F_2N_2O_3Si$ $[M]^+$ *m/z* 575.2542, found 575.2616.

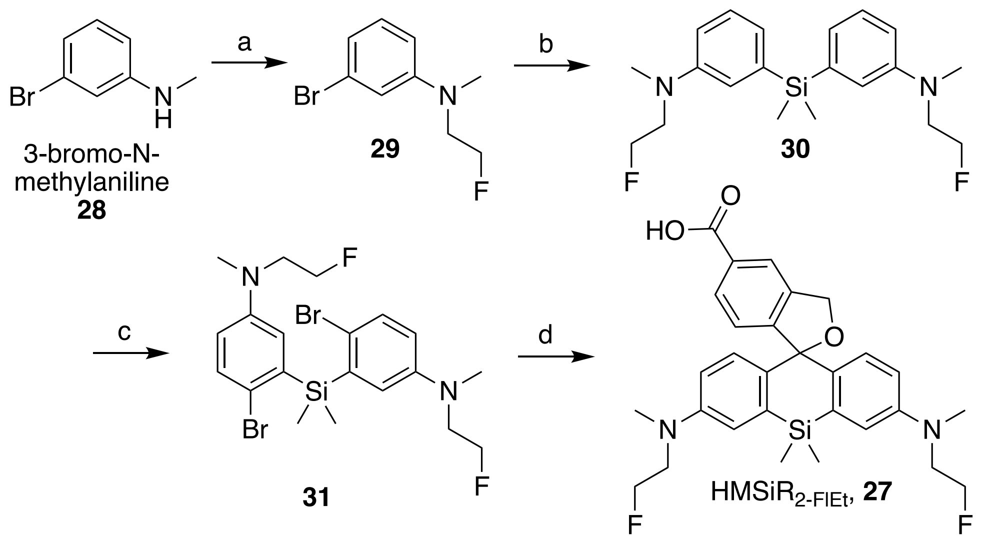

**H. Supplementary Scheme S6.** Synthesis of HMSiR$_{2Fl\text{-}Et}$.

a) 1-fluoro-2-iodoethane, $K_2CO_3$, DMF, 70˚C, b) n-BuLi, $Me_2SiCl_2$, -78 ˚C to rt, c) NBS, DMF, rt, d) i. tert-BuLi, **3**, -78 ˚C to rt, ii. TFA, rt

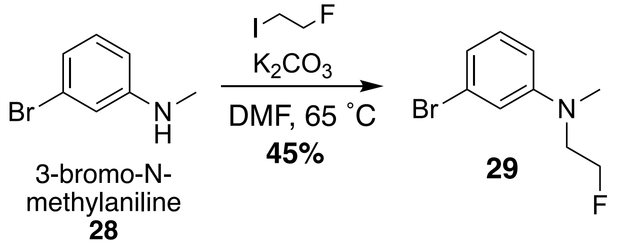

**3-bromo-*N*-(2-fluoroethyl)-*N*-methylaniline 29.** Potassium carbonate (1.1 g, 8.06 mmol, 1.5 equiv) was added to a solution of 3-bromo-N-methylaniline **28** (1.0 g, 5.38 mmol, 1.0 equiv) in DMF (4.8 mL). 1-bromo-2-fluoroethane (0.53 mL, 6.45 mmol, 1.2 equiv) was added, and the reaction was stirred at 65 °C for 18 h. The mixture was allowed to cool to room temperature, diluted with water, and acidified with 1 M HCl. The aqueous phase was extracted with ethyl acetate, and the combined organic phases were washed 3x with water and brine and dried over $Na_2SO_4$. The solvent was evaporated under vacuum, and the crude product was purified by flash column chromatography on silica (100% hexanes to EtOAc:hexanes 5:95) to yield **29** (53%) as a dark oil. $^1$H-NMR ($CDCl_3$, 400 MHz) δ 7.06 (t, *J* = 7.8 Hz, 1 H), 6.82 (m, 2 H), 6.60 (m, 1 H), 4.63 (t, *J* = 5.2 Hz, 1 H), 4.51 (t, *J* = 5.2 Hz, 1 H), 3.64 (t, *J* = 5.2 H, 1 H), 2.98 (s, 3 H). $^{13}$C NMR (101 MHz, $CDCl_3$) δ 150.17, 130.58, 123.68, 119.74, 115.19, 110.98, 82.58, 80.90, 77.48, 77.16, 76.84, 52.83, 52.62, 39.21.

**3,3'-(dimethylsilanediyl)bis(*N*-(2-fluoroethyl)-*N*-methylaniline) 30.** Aniline **29** (478 mg, 2.06 mmol, 2.4 equiv) was dissolved in THF (4.5 mL) in a flame-dried round bottom flask and cooled

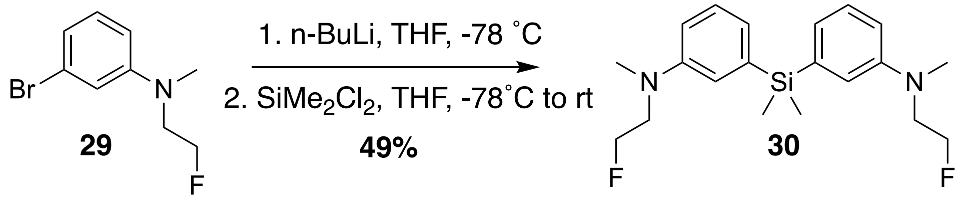


to -78 °C under nitrogen. *n*-Butyllithium (2.5 M in cyclohexane, 0.82 mL, 2.06 mmol, 2.4 eq) was added, and the reaction was stirred at -78 °C for 30 min. Dichlorodimethylsilane (0.11 mL, 0.86 mmol) in THF (0.5 mL) was then added, the dry ice bath was removed, and the reaction was stirred at room temperature for 3 h. It was subsequently quenched with saturated $NH_4Cl$, diluted with water, and extracted with EtOAc (2×). The combined organic phase was washed with brine, dried over anhydrous $Na_2SO4$, filtered, and concentrated under vacuum. The crude product was purified by flash chromatography on silica (100% hexanes to EtOAc:hexanes 15:85) to yield **30** as a golden oil (49%). $^1$H-NMR ($CDCl_3$, 400 MHz) δ 7.24 (m, 2 H), 6.90 (m, 4 H), 6.75 (d, J = 8.6 Hz, 2 Hz), 4.64 (t, *J* = 6.0 Hz, 2 H), 4.52 (t, *J* = 6.0 Hz, 2 H), 3.67 (t, *J* = 5.2 Hz, 2 H), 3.50 (t, 5.2 Hz, 2 H), 3.0 (s, 6 H), 0.53 (s, 6 H). $^{13}$C NMR (101 MHz, $CDCl_3$) δ 148.26, 139.24, 128.77, 122.91, 117.99, 113.33, 82.71, 81.02, 77.42, 77.10, 76.79, 53.00, 52.79, 39.12, -2.20.

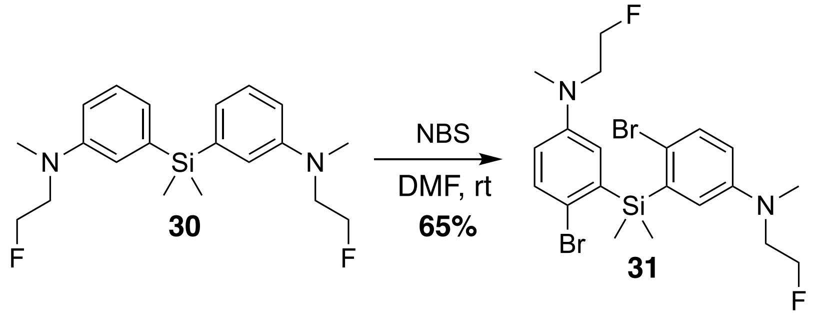


**3,3'-(dimethylsilanediyl)bis(4-bromo-*N*-(2-fluoroethyl)-*N*-methylaniline) 31.** Silane **30** (105.8 mg, 0.28 mmol) was taken up in DMF (1.0 mL). N-Bromosuccinimide (103.89 mg, 0.58 mmol, 2.0 eq) was dissolved in 1.0 mL DMF and added dropwise over 5 min at 0 ˚C. The reaction was then stirred at room temperature overnight. Product formation was confirmed by LCMS, and then the resulting residue was concentrated in vacuo, diluted with water, and extracted twice with EtOAc.

The combined organic extracts were washed with water and brine, dried over anhydrous MgSO4, filtered, and concentrated in vacuo. Silica gel chromatography (100% hexanes to EtOAc:hexanes 15:85) afforded **31** as a reddish brown solid (65%). $^1$H-NMR ($CDCl_3$, 400 MHz) δ 7.40 (d, *J* = 8.8 Hz, 2 H), 6.82 (d, J = 3.3 Hz, 2 H), 6.65 (dd, J = 8.9, 3.3 Hz, 2 H), 4.63 (t, *J* = 5.3 Hz, 2 H), 4.51 (*J* = 5.3 Hz, 2 H), 3.62 (t, *J* = 5.2 Hz, 2 H), 3.56 (t, *J* = 5.2 Hz, 2H), 2.99 (s, 6 H), 0.79 (s, 6 H). $^{13}$C NMR (101 MHz, $CDCl_3$) δ 147.44, 139.05, 133.33, 121.60, 117.01, 115.22, 82.68, 80.99, 77.48, 77.16, 76.84, 53.15, 52.95, 39.20, -0.79.

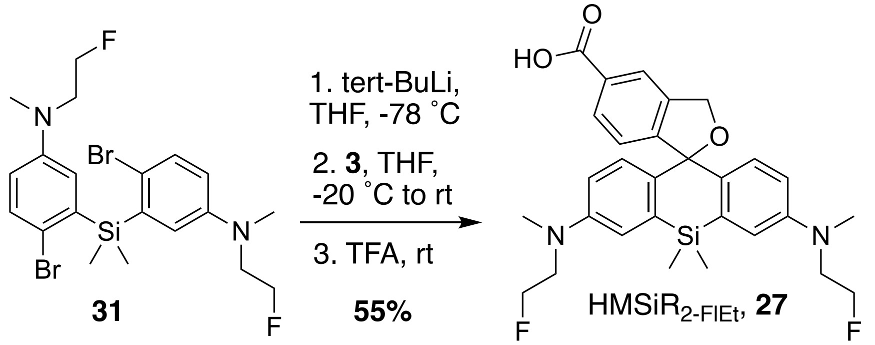


**HMSiR$_{2\text{-FlEt}}$ 27.** A solution of dibromide **31** (48.0 mg, 0.092 mmol) in THF (1.6 mL) was cooled to -78 °C under nitrogen. *tert*-Butyllithium (1.7 M in pentane, 0.24 mL, 0.406 mmol, 4.4 eq) was added, and the reaction was stirred at -78 °C for 30 min. A solution of **31** (47.5 mg, 0.203 mmol, 2.2 eq) in THF (1.6 mL) was added dropwise over 30 min. The reaction was allowed to warm to room temperature overnight. Product formation was confirmed by LCMS, and it was subsequently diluted with saturated NH4Cl and water and extracted with EtOAc (2×). The combined organic extracts were washed with brine, dried over anhydrous MgSO4, filtered, and concentrated *in vacuo*. The resulting residue was redissolved in MeOH (3 mL), and AcOH (100 µL) was added. After stirring for 10 min, the solution was concentrated to dryness and purified by flash chromatography on silica (100% $CH_2Cl_2$ to $CH_2Cl_2$:MeOH 90:10). The resultant material was stirred in TFA for 30 min, concentrated *in vacuo* and re-purified by column chromatography (100% $CH_2Cl_2$ to $CH_2Cl_2$:MeOH 90:10) to afford **HMSiR$_{2\text{FlEt}}$** as a bluish green solid (55%). $^1$H-NMR (MeOD, 400 MHz) δ 7.97 (s, 1 H), 7.80 (d, *J* = 7.8 Hz, 1 H), 7.04 (d, *J* = 8.7 Hz, 2 H), 6.98 (d, *J* = 2.9 Hz, 2 H), 6.89 (d, *J* = 7.9 Hz, 1 H), 6.64 (dd, J = 9.0, 2.9 Hz, 2 H), 5.41 (s, 2 H), 4.60 (t, J = 5.1 Hz, 2 H), 4.49 (t, J = 5.1 Hz, 2 H), 3.66 (t, J = 5.1 Hz, 2 H), 3.61 (t, J = 5.1 Hz, 2 H) 2.95 (s, 6 H), 0.55 (s, 3 H), 0.42 (s, 3 H). HRMS calc'd for $C_{29}H_{33}F_2N_2O_3Si$ $[M]^+$ *m/z* 523.2229, found 523.2320.

**I. Supplementary Scheme S7.** Synthesis of $Cal_{664sb}$.

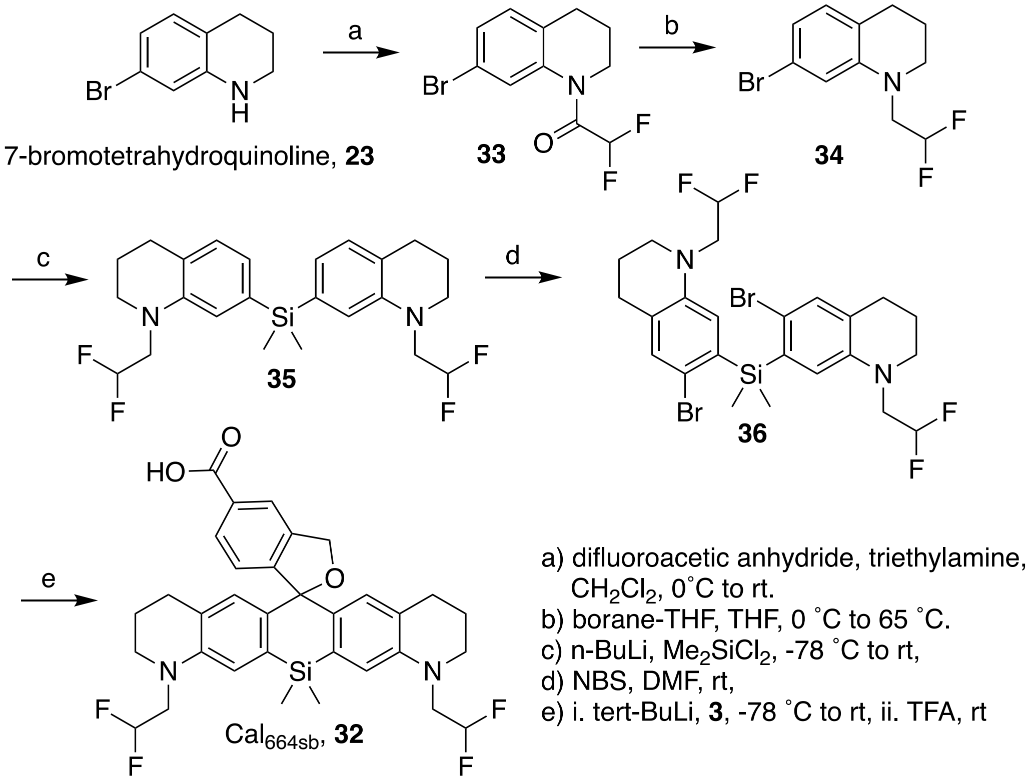


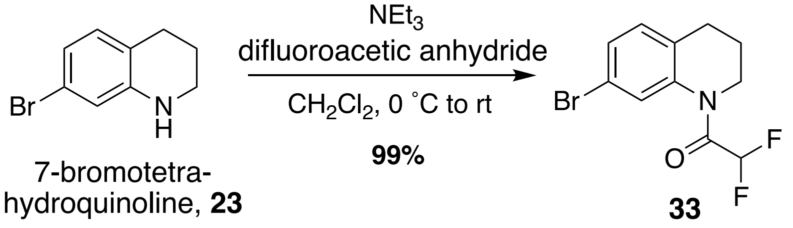


**1-(7-bromo-3,4-dihydroquinolin-1(2*H*)-yl)-2,2-difluoroethan-1-one 33.** 7-bromotetrahydroquinoline **23** (318.1 mg, 1.5 mmol) was dissolved in $CH_2Cl_2$ (10 mL) and $Et_3N$ (0.44 mL, 3.15 mmol, 2.1 eq) was added. An aliquot of $(CF_2CO)_2O$ (0.23 mL, 1.8 mmol, 1.2 eq) was added dropwise at 0 °C, and the mixture was stirred for 24 h at room temperature. The reaction mixture was washed with saturated $NaHCO_3$(aq). The organic layer was separated, and the aqueous layer was back-extracted twice with $CH_2Cl_2$. The combined organic extracts were dried over $Na_2SO_4$ and evaporated to yield **33** as a light yellow solid (430 mg, 99%). The product was used in the following synthetic step without further purification.

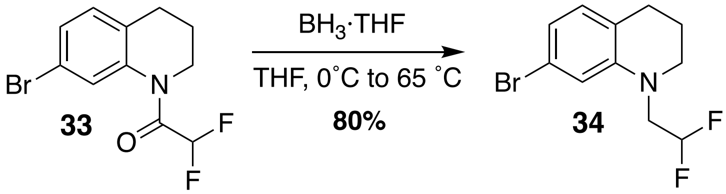

**7-bromo-1-(2,2-difluoroethyl)-1,2,3,4-tetrahydroquinoline 34.** To a solution of **33** (290.1 mg, 1.0 mmol) in dry THF (1 mL) at 0°C was added an aliquot of 1M solution $BH_3$ in THF (2 mL, 2.0 eq). The mixture was then heated at reflux for 16 hr. The reaction mixture was cooled to 0°C and the excess $BH_3$ carefully quenched upon addition of MeOH (1 mL) followed by 1 M aq. NaOH (1 mL). After stirring at room temperature for 20 min, the mixture was diluted with ether (10 mL), and the organic layer was separated. The aqueous layer was extracted with ether (3×10 mL); combined organic solutions were washed with sat. aq. $NaHCO_3$, brine, dried and the solvent was evaporated in vacuo. The desired product was purified by flash column chromatography (220.0 mg, 80%)

**bis(1-(2,2-difluoroethyl)-1,2,3,4-tetrahydroquinolin-7-yl)dimethylsilane 35.** 7-bromo-1-(2,2-difluoroethyl)-1,2,3,4-tetrahydroquinoline **34** (198.8 mg, 0.72 mmol, 2.4 eq) was dissolved in 1.5 mL THF in a flame-dried round-bottom flask and cooled to -78 °C under nitrogen. *n*-Butyllithium (0.29 mL, 2.5 M in cyclohexane, 2.4 eq) was added, and the reaction was stirred at -78 °C for 30 min. Dichlorodimethylsilane (36.5 uL, 0.3 mmol, dissolved in 0.2 mL THF) was then added, the dry ice bath was removed, and the reaction was stirred at room temperature for 3 h. It was subsequently quenched with saturated $NH_4Cl$, diluted with water, and extracted with EtOAc (2×10 mL). The combined organic phase was washed with brine, dried over anhydrous $Na_2SO4$, filtered, and concentrated under vacuum. The crude product was purified by flash chromatography on silica (100% hexanes to EtOAc:hexanes 15:85) to yield **35** as a colorless oil (83.9 mg, 62%).

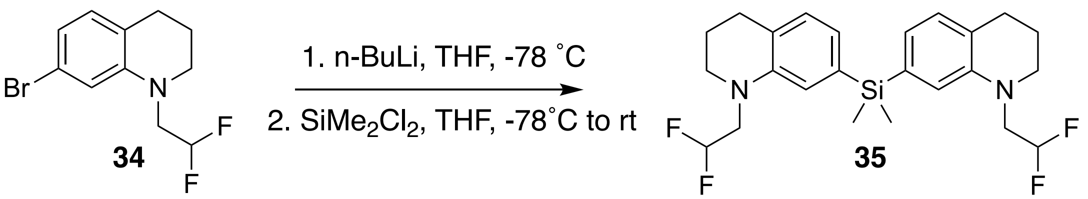

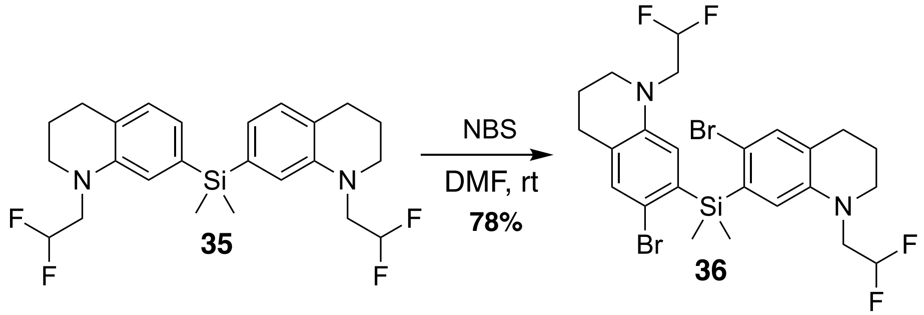


**bis(6-bromo-1-(2,2-difluoroethyl)-1,2,3,4-tetrahydroquinolin-7-yl)dimethylsilane 36.** Silane **35** (120.1 mg, 0.267 mmol) was taken up in 2 mL DMF. N-Bromosuccinimide (94.9 mg, 0.533 mmol, 2.0 eq) was dissolved in 1 mL DMF and added dropwise over 5 min at 0 ˚C. The reaction was then stirred at room temperature overnight. Product formation was confirmed by LCMS, and then the resulting residue was concentrated in vacuo, diluted with water and extracted twice with EtOAc (20 mL). The combined organic extracts were washed with water and brine, dried over anhydrous $MgSO_4$, filtered, and concentrated in vacuo. Silica gel chromatography (100% hexanes to EtOAc:hexanes 15:85) afforded **26** as a white solid (126.7 mg, 78%)

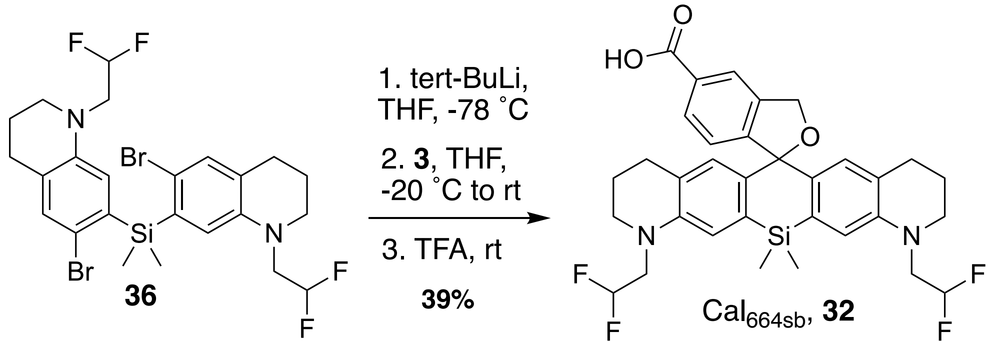


**Cal$_{664sb}$, 32**. A solution of dibromide **36** (41.5 mg, 0.068 mmol) in THF was cooled to -78 °C under nitrogen. *tert*-Butyllithium (0.18 mL, 0.30 mmol, 1.7 M in pentane, 4.4 eq) was added, and the reaction was stirred at -78 °C for 30 min. A solution of **3** (35.2 mg, 0.15 mmol, 2.2 eq) in 1.5 mL THF was added dropwise over 30 min. The reaction was allowed to warm to room temperature overnight. Product formation was confirmed by LCMS. The reaction was subsequently diluted with saturated $NH_4Cl$ and water and extracted twice with EtOAc. The combined organic extracts were washed with brine, dried over anhydrous $MgSO_4$, filtered, and concentrated *in vacuo*. The resulting residue was redissolved in MeOH (3 mL), and AcOH (100 µL) was added. After stirring for 10 min, the solution was concentrated to dryness and purified by flash chromatography on silica (100% $CH_2Cl_2$ to $CH_2Cl_2$:MeOH 90:10). The resultant material was stirred in TFA for 30 min, concentrated *in vacuo* and re-purified by column chromatography (100% $CH_2Cl_2$ to $CH_2Cl_2$:MeOH 90:10) to afford Cal$_{664sb}$ as a bright blue solid (16.2 mg, 39% over two steps).

**J. Supplementary Scheme S8.** Synthesis of Yale$_{676sb}$-CA.

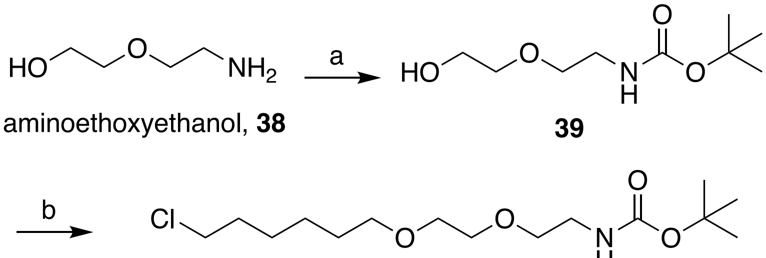

aminoethoxyethanol, **38**

**39**

**40**

a) $Boc_2O$, EtOH, rt, b) i. NaH, THF/DMF, 0 ˚C, ii. 1-chloro-6-iodohexane, THF/DMF, 0 ˚C to rt, c) TFA, $CH_2Cl_2$, 0˚C to rt

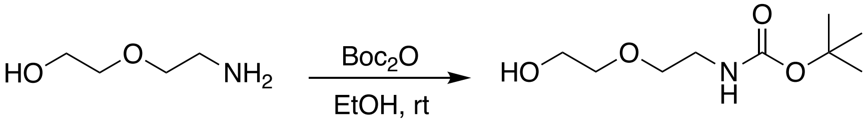

aminoethoxyethanol, **38** **99%** **39**

***tert*-butyl (2-(2-hydroxyethoxy)ethyl)carbamate 39.** To a solution of 2-(2-aminoethoxy)ethanol (2.0 g, 19.02 mmol) in anhydrous EtOH (40 mL) was added $Boc_2O$ (4.36 g, 19.02 mmol) at 0 °C. After stirring at room temperature 2 h, the reaction mixture was evaporated. The product was then extracted thrice with $CH_2Cl_2$. The combined organic layers were dried over $Na_2SO_4$ and evaporated under vacuum to obtain the product as colorless oil (99%), which was used for the next step without further purification. $^{1H}$NMR: (400 MHz, $CDCl_3$, δ): 5.28 (br, 1H), 3.68-3.52 (m, 2H), 3.52-3.48 (m, 4H), 3.28-3.26 (m, 2H), 1.39 (s, 9H). $^{13}$CNMR: (100MHz, CDCl3, δ): 156.32, 79.41, 72.37, 70.37, 61.62, 40.42, 28.48. HRMS (ESI, m/z): [M+Na] calcd for $C_9H_{19}NNaO_4$, 228.1212; found 228.1207.

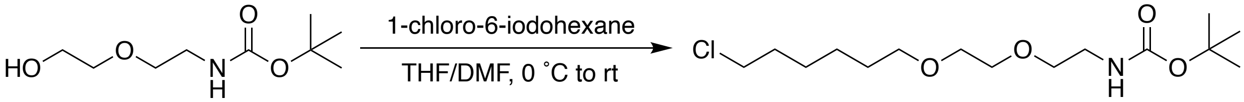

**39** **69%** **40**

***tert*-butyl (2-(2-((6-chlorohexyl)oxy)ethoxy)ethyl)carbamate.** To a solution of **39** (1.0 g, 4.88 mmol) in THF (7 mL) and DMF (3.5 mL) at 0 °C was added NaH (233.78 mg, 60% in mineral oil, 5.85 mmol). After stirring at 0 °C for 30 min, 6-chloro-1-iodohexane (1.0 mL, 6.82 mmol) was added to the above solution. The reaction mixture was stirred overnight and quenched with saturated NH4Cl. The mixture was extracted with EtOAc (3x), washed with $H_2O$ and brine. The combined organic layers were dried over $Na_2SO_4$ and concentrated. The crude product was purified by silica gel column chromatography (100% hexanes to 35:65 EtOAc:Hexanes) to yield pure **40** (69%) as colorless oil. 1H NMR: (400MHz, CDCl3, δ): 5.03 (br, 1H), 3.59-3.57 (m, 2H), 3.55-3.49 (m, 6H), 3.43 (t, $J$ = 6.6 Hz,, 2H), 3.31-3.27 (m, 2H), 1.79-1.72 (m, 2H), 1.62-1.55 (m, 2H), 1.47-1.33 (m, 13H); 13CNMR: (100MHz, CDCl3, δ): 156.13, 79.28, 71.39, 70.36, 70.32,

70.14, 45.16, 40.45, 32.65, + 29.54, 28.54, 26.80, 25.53; HRMS (ESI, m/z): [M+Na] calcd for C15H30ClNNaO4, 346.1761; found 346.1752.

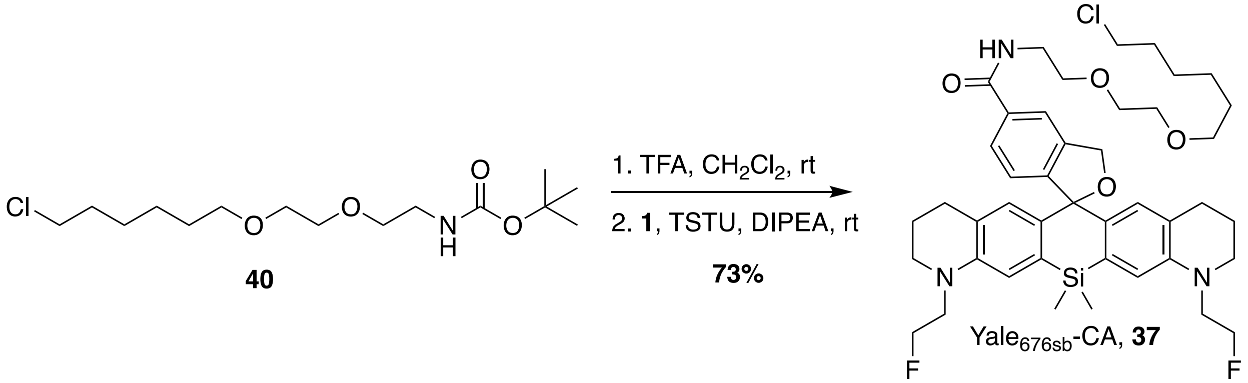


**Yale$_{676sb}$-CA, 37.** To a solution of **40** in $CH_2Cl_2$ was added TFA. The solution was stirred at room temperature for 1 hr and the progress of the reaction was monitored by TLC. After completion of the reaction, the reaction mixture was concentrated *in vacuo*. To the crude residue was added a pre-mixed solution (for 15 min) of **1**, TSTU and DIPEA. Additional DIPEA was added and then the reaction was stirred at room temperature for 3 h. Following completion of the reaction, as monitored by LCMS, the product was purified *via* reverse-phase HPLC to yield Yale$_{676sb}$-CA as a blue solid.

### II. Biophysical experiments

**A. Absorption and fluorescence spectra.** Solutions used for absorption and emission spectra were obtained by diluting a 5 mM stock solution of each fluorophore with 0.2 M aqueous sodium phosphate buffer to a final concentration of 0.5 μM. Absorption spectra were measured at room temperature using a Beckman Coulter DU 730 UV/Vis spectrophotometer and 1 cm pathlength, 0.1 mL microcuvettes. Fluorescence spectra were measured at room temperature using a TCSPC TD-Fluor Horiba Fluorolog 3 Time Domain Fluorimeter using 1 cm pathlength, 3.5 mL cuvettes. All absorption and fluorescence spectra were acquired at a pH that maximized the concentration of the open/emissive form of each fluorophore (pH = 2.0 for HMSiR$_{indol}$, HMSiR$_{julol}$, HMSiR$_{THQ}$ and Yale$_{676sb}$; pH = 2.5 for HMSiR$_{2\text{-FlEt}}$; pH = 4.5 for HMSiR).

**B. Plot of Quantum Yield *vs.* Absorption Maxima for Rhodamine-derived Fluorophores**

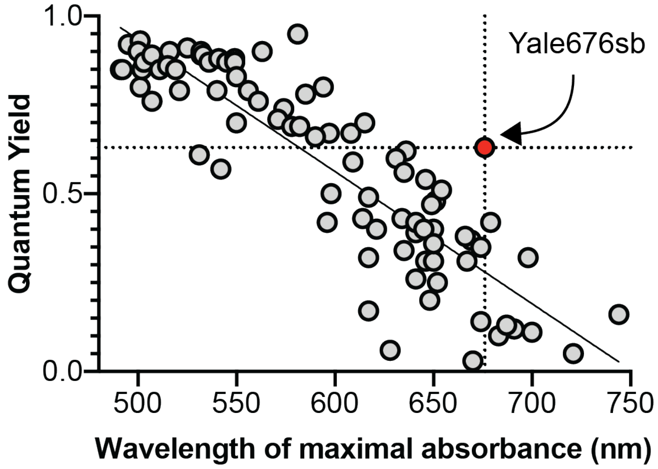


**Supplementary Figure 1.** Plot of quantum yield as a function of the wavelength of maximal absorbance ($\lambda_{abs}$) for previously reported rhodamine-derived fluorophores and $Yale_{676sb}$.

**C. Table of Quantum Yield and Absorption Maxima for Rhodamine-derived Fluorophores**

**Supplementary Table 1.**

| Name: | Type | $\lambda_{abs}$ (nm) | $\lambda_{em}$ (nm) | Quantum Yield | Reference |
|---|---|---|---|---|---|
| 2-Me TokyoGreen | fluorescein | 491 | 510 | 0.85 | 1 |
| Fluorescein | fluorescein | 492 | 511 | 0.85 | 2 |
| 500R | rhodamine | 501 | 523 | 0.93 | 3 |
| HMRG | rhodamine | 500 | 523 | 0.9 | 4 |
| HMbisTFER | rhodamine | 502 | 526 | 0.85 | 4 |
| JF503 | rhodol | 503 | 529 | 0.87 | 5 |

| | | | | | |
|---|---|---|---|---|---|
| HEtetTFER | rhodamine | 507 | 530 | 0.76 | 4 |
| HMtetTFER | rhodamine | 507 | 532 | 0.89 | 4 |
| 520R | rhodamine | 521 | 543 | 0.79 | 3 |
| 515R | rhodamine | 515 | 543 | 0.86 | 3 |
| JF519 | rhodol | 519 | 546 | 0.85 | 5 |
| JF525 | rhodamine | 525 | 549 | 0.91 | 5 |
| 530RH | rhodamine | 532 | 553 | 0.89 | 6 |
| Molecule #11 | rhodamine | 533 | 557 | 0.89 | 5 |
| Molecule #10 | rhodamine | 536 | 560 | 0.87 | 5 |
| Molecule #9 | rhodamine | 541 | 564 | 0.88 | 5 |
| Molecule #8 | rhodamine | 542 | 565 | 0.57 | 5 |
| Molecule #7 | rhodamine | 545 | 568 | 0.87 | 5 |
| JF549 | rhodamine | 549 | 571 | 0.88 | 5 |
| Molecule #6 | rhodamine | 549 | 572 | 0.87 | 5 |
| Molecule #5 | rhodamine | 550 | 572 | 0.83 | 5 |
| TMR | rhodamine | 550 | 574 | 0.7 | 7 |
| 560CP | C-rhodamine | 561 | 588 | 0.76 | 6 |
| 575RH | rhodamine | 574 | 597 | 0.74 | 6 |
| 570CPH | C-rhodamine | 571 | 600 | 0.71 | 6 |
| 580CP | C-rhodamine | 582 | 607 | 0.69 | 3 |

| | | | | | |
|---|---|---|---|---|---|
| 580R | rhodamine | 581 | 607 | 0.95 | 3 |
| JF585 | C-rhodamine | 585 | 609 | 0.78 | 5 |
| Maryland Red | Si-fluorescein | 597 | 613 | 0.67 | 8 |
| Molecule #26 | Si-fluorescein | 598 | 614 | 0.5 | 8 |
| Molecule #24 | Si-fluorescein | 596 | 618 | 0.42 | 8 |
| SnP | pyronine | 614 | 628 | 0.43 | 9 |
| JF608 | C-rhodamine | 608 | 631 | 0.67 | 5 |
| 610CP | C-rhodamine | 609 | 634 | 0.59 | 10 |
| GeP | pyronine | 621 | 634 | 0.4 | 9 |
| Molecule #27 | Si-fluorescein | 617 | 635 | 0.32 | 8 |
| 620SiR | Si-rhodamine | 617 | 638 | 0.49 | 10 |
| 620CP | C-rhodamine | 617 | 647 | 0.17 | 10 |
| Molecule #50 | pyronine | 636 | 649 | 0.62 | 8 |
| GeR | Ge-rhodamine | 635 | 649 | 0.34 | 9 |
| 630GeRH | Ge-rhodamine | 631 | 651 | 0.6 | 6 |
| JF635 | Si-rhodamine | 635 | 652 | 0.56 | 5 |
| GeR | X-rhodamine | 634 | 655 | 0.43 | 6 |
| Molecule #51 | pyronine | 641 | 657 | 0.26 | 8 |
| TMDHS | pyronine | 641 | 659 | 0.39 | 11 |
| 630CP | C-rhodamine | 628 | 660 | 0.06 | 3 |
| SiR-Me | Si-rhodamine | 646 | 660 | 0.31 | 9 |

| | | | | | |
|---|---|---|---|---|---|
| Molecule #44 | rosamine | 648 | 662 | 0.2 | 8 |
| Molecule #48 | Si-rhodamine | 651 | 666 | 0.48 | 8 |
| 640SiRH | Si-rhodamine | 641 | 662 | 0.42 | 6 |
| Molecule #45 | Si-rhodamine | 649 | 663 | 0.47 | 8 |
| JF646 | Si-rhodamine | 646 | 664 | 0.54 | 5 |
| Molecule #41 | Si-rhodamine | 650 | 667 | 0.4 | 8 |
| carboxy-SiR | Si-rhodamine | 645 | 667 | 0.4 | 7 |
| Molecule #43 | Si-rhodamine | 652 | 668 | 0.25 | 8 |
| Molecule #42 | Si-rhodamine | 654 | 670 | 0.51 | 8 |
| HMSiR | Si-rhodamine | 650 | 671 | 0.31 | 4 |
| 650SiR | Si-rhodamine | 650 | 672 | 0.36 | 3 |
| JF669 | Si-rhodamine | 669 | 682 | 0.37 | 8 |
| Molecule #37 | Si-rhodamine | 667 | 682 | 0.31 | 8 |
| Molecule #39 | Si-rhodamine | 674 | 685 | 0.14 | 8 |
| NR666 | X-rhodamine | 666 | 685 | 0.38 | 12 |
| SiR680 | Si-rhodamine | 674 | 689 | 0.35 | 13 |
| 670SiR | Si-rhodamine | 670 | 696 | 0.03 | 3 |
| 680SiR | Si-rhodamine | 679 | 697 | 0.42 | 10 |
| Molecule #38 | Si-rhodamine | 683 | 698 | 0.1 | 8 |
| SiR700-Me | Si-rhodamine | 691 | 712 | 0.12 | 13 |

| | | | | | |
|---|---|---|---|---|---|
| NR698 | X-rhodamine | 698 | 712 | 0.32 | 12 |
| carboxy-SiR700 | Si-rhodamine | 687 | 716 | 0.13 | 7 |
| NR700 | X-rhodamine | 700 | 722 | 0.11 | 12 |
| SiR720 | Si-rhodamine | 721 | 740 | 0.05 | 13 |
| NR744 | X-rhodamine | 744 | 764 | 0.16 | 12 |

**D. Determination of p$K_{cycl}$.** The p$K_{cycl}$ of each fluorophore was determined from a plot of the absorbance of the open/emissive form as a function of pH. Samples of each fluorophore at 2 μM in 0.2 M sodium phosphate were prepared at half-integral pH values between 2 and 12 and in triplicate using a 96-well plate. The absorbance of each solution at each pH at the wavelength corresponding to the emission $\lambda_{max}$ was measured at room temperature. The data was plotted using GraphPad Prism 8.0. The normalized curves for fluorophores with a single titratable group in this pH range (Yale$_{676sb}$, HMSiR$_{indol}$, HMSiR$_{julol}$, and Cal$_{664sb}$) were fit using the following equation:

$$abs = (c_0 + c_1 * 10^{pH - pKcycl})/(1 + 10^{pH - pKcycl})$$

The normalized curves for fluorophores with two titratable groups in this pH range (HMSiR and HMSiR$_{2\text{-FlEt}}$) were fit using the following equation:

$$abs = (c_0 + c_1 * 10^{pH - pKcycl1} + c_2 * 10^{2*pH - pKcycl1 - pKcycl2 \blacksquare})/(1 + 10^{pH - pKcycl})$$

All curves were fit such that the value of p$K_{cycl}$, p$K_{cycl1}$ and p$K_{cyl2}$ were set to the value of pH at which the normalized absorbance = 0.5. For fluorophores with both p$K_{cycl1}$ and p$K_{cycl2}$ (HMSiR and HMSiR$_{2\text{-FlEt}}$), p$K_{cycl2}$ was arbitrarily set as greater than p$K_{cycl1}$, and p$K_{cycl2}$ was reported.

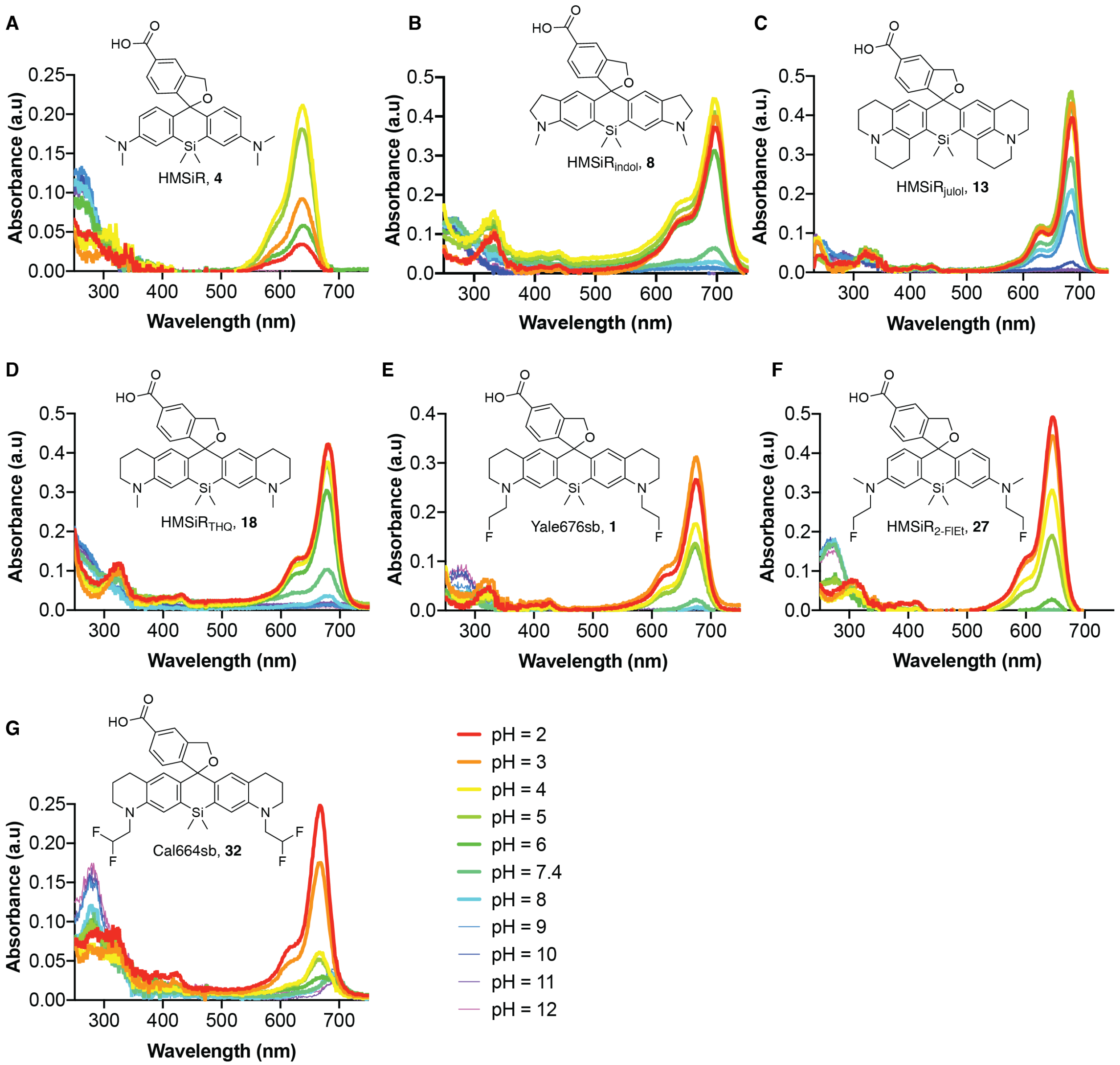


**Supplementary Figure 2**. pH-dependent absorption spectra of (a) HMSiR, (b) $HMSiR_{indol}$, (c) $HMSiR_{julol}$, (d) $HMSiR_{THQ}$, (e) $Yale_{676sb}$, (f) $HMSiR_{2\text{-}FlEt}$, and (g) $Cal_{664sb}$ at concentrations between 2 to 5 μM and at various pH values in 0.2 M sodium phosphate buffer containing 0.1% DMSO as a cosolvent.

E. **Determination of Quantum Yield.** Absolute fluorescence quantum yield values were measured under identical conditions using a Quantaurus-QY Plus spectrometer (model C13534, Hamamatsu). This instrument uses an integrating sphere to determine photons absorbed and emitted by a sample. Measurements were carried out using 5 μM samples in PBS +0.01% DMSO buffered at pH = 4.5 (HMSiR) or pH = 2.5 (HMSiR$_{indol}$, HMSiR$_{julol}$, HMSiR$_{THQ}$, Yale$_{676sb}$, HMSiR$_{2\text{-}FlEt}$, and Cal$_{664sb}$). Reported values are averages (n = 3).

**F. Single-molecule kinetics measurements.** Kinetic data were acquired using dye molecules immobilized on glass and imaged with a variety of laser powers at an acquisition rate of 400 Hz. During analysis, consecutive frames over which a molecule was detected were grouped into ‘emissions’. On-times were determined from the duration of emissions. Off-times correspond to the number of frames between two consecutive emissions of the same molecule. Numbers of switching cycles were determined by the number of emissions throughout the acquisition, over which all molecules were bleached.

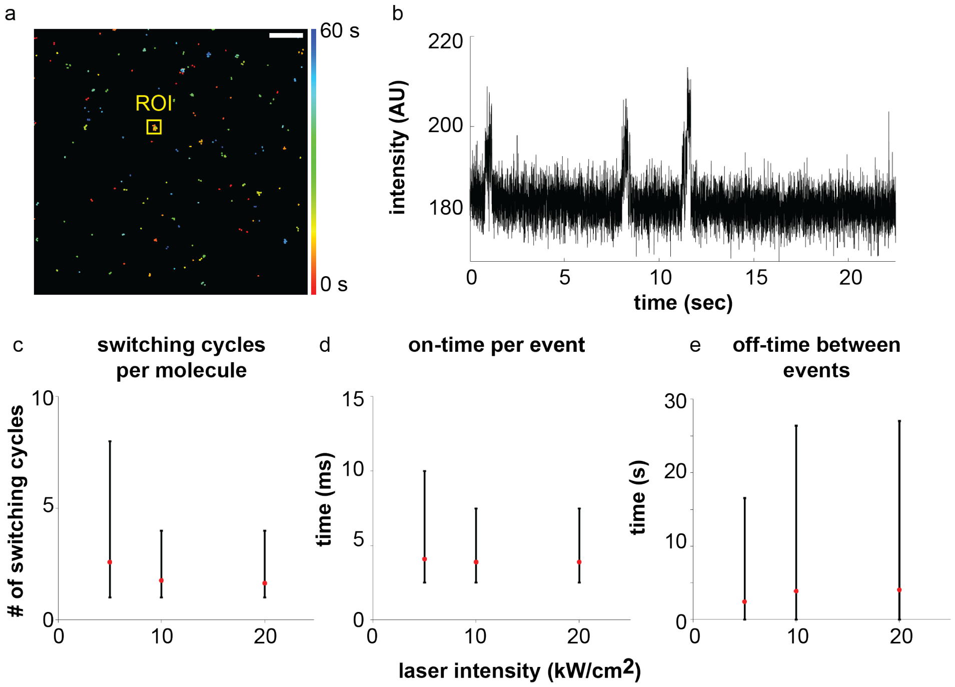


**Supplementary Figure 4.** (A) Super-resolved image of Yale$_{676sb}$ immobilized on glass colored by time of emission. Intensity trace (B) of single molecule shown in ROI of (A). Plots showing mean, 5th, and 95th percentiles for number of switching cycles (C), on-times (D), and off-times (E) for single molecules. Scale bar 1 μm. All kinetics data used 120 s of acquired frames.

### III. Microscopy

A. **Cross-talk measurements.** Cross-talk was determined using single-color control samples labeled, imaged, and analyzed as in the two-color imaging. Under these conditions, $Yale_{676sb}$ exhibited 6.4% crosstalk at a 13% rejection rate while HMSiR had 5.6% crosstalk at a 20% rejection rate (n = 3 for $Yale_{676sb}$, n = 5 for HMSiR, with at least 1 million localizations per acquisition).

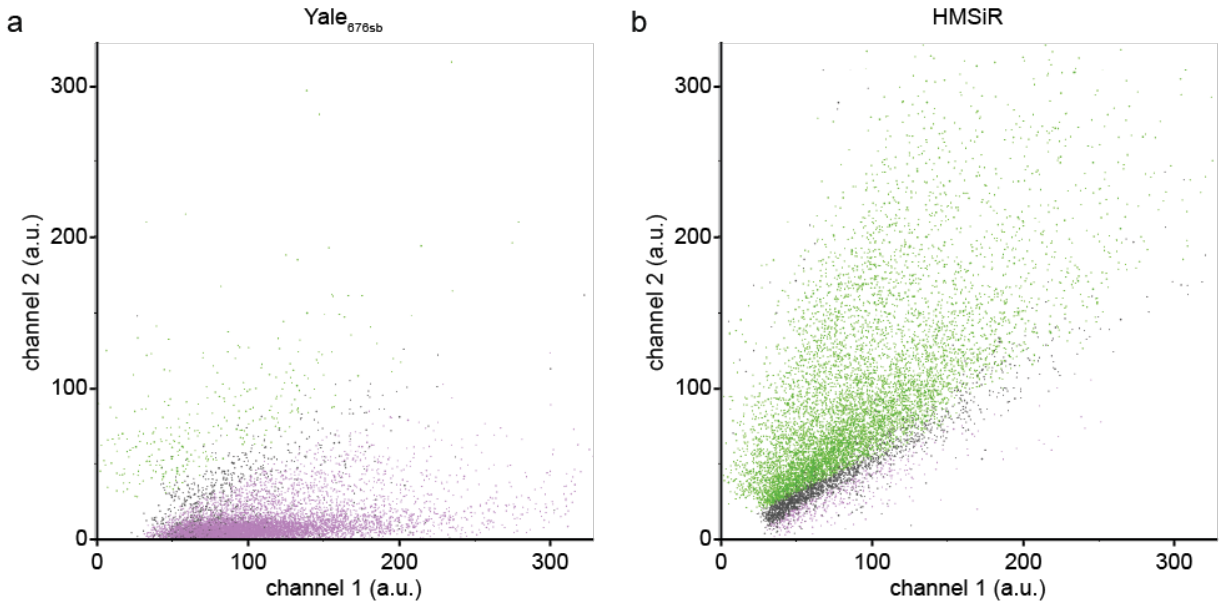


**Supplementary Figure 5.** Intensity in two camera channels for single-molecules in representative $Yale_{676sb}$ (a) and HMSiR (b) one-color samples. Molecules highlighted in purple are assigned to $Yale_{676sb}$ while those in green are assigned to HMSiR and those in black are rejected.

**B. Labeling Halo-and-SNAP-fusion proteins.** U-2 OS HTB-96 cells (ATCC) were electroporated with Halo-Sec61β (and SNAP-OMP25) for two-color samples) then seeded into a mattek dish (MatTek Corporation, P35G-1.5-20-C).[14] After incubating overnight, the cells were live labeled with 300nM Yale676sb-CA (and 300nM HMSiR-BG for two-color samples) diluted in McCoy's 5A (ThermoFisher Scientific, 12330031) with 1% casein for 1 hour. Cells were rinsed three times and then the media was changed once more after 30 minutes. The cells were imaged in Live Cell Imaging Solution (ThermoFisher Scientific, A14291DJ) that was supplemented with 15mM glucose. Prior to seeding, Mattek dishes were sonicated for 15 minutes in 1M KOH, rinsed three times with Milli-Q water, sterilized with 100% ethanol, and then air dried in a biological safety cabinet.

**C. Super-resolution microscopy and image analysis.** 2D super-resolution images were recorded using a custom-built ratiometric setup running custom LabVIEW software. A 642 nm laser (2W; MPB Communications) was used to excite Yale676 and HMSiR. Laser power was controlled using an acousto-optic tunable filter (AOTFnC-400.650-TN; AA Opto Electronic). Fluorescence was collected using an oil immersion objective lens (63x, 1.47 NA; Leica), passed through a quad-band filter (ZET405/488/561/647m; Chroma) and a bandpass filter (FF01-709/167; Semrock) before being imaged on a sCMOS camera (Orca Flash 4.0; HamamatsuPhotonics). For two-color imaging, channels were split by a bandpass filter (FF01-694/SP; Semrock) and the transmitted channel additionally passed through another bandpass filter (FF01-670/30; Semrock). A pair of relay lenses was used to achieve an effective pixel size of 99 nm. Images were acquired at 400 frames per second under even wide-field illumination of 10 kW/cm$^2$ with a 40 µm square field of view. A custom-built focus lock system was used to prevent axial drift of the sample during data acquisition. A 850 *nm* fiber-coupled diode laser (L850P010; Thorlabs) was introduced using total internal reflection and its reflection from the coverglass-media interface was imaged onto a USB CMOS camera (DCC1545M; Thorlabs). The position of the reflected light was localized and used as feedback to adjust a piezoelectric objective positioner (P726; Physik Instrumente). Samples were incubated at 37 C using a stagetop incubator system (STXF-WSKMX-SET; Tokai Hit). All localization, visualization, and image analysis were performed using PYthon Microscopy Environment (PYME).

(4) Uno, S.; Kamiya, M.; Morozumi, A.; Urano, Y. A Green-Light-Emitting, Spontaneously Blinking Fluorophore Based on Intramolecular Spirocyclization for Dual-Colour Super-

Resolution Imaging. *Chem. Commun.* **2017**, *54* (1), 102–105. https://doi.org/10.1039/c7cc07783a.

https://doi.org/10.1021/ja210375e.

(14) Bottanelli, F.; Kromann, E. B.; Allgeyer, E. S.; Erdmann, R. S.; Wood Baguley, S.; Sirinakis, G.; Schepartz, A.; Baddeley, D.; Toomre, D. K.; Rothman, J. E.; Bewersdorf, J. Two-Colour Live-Cell Nanoscale Imaging of Intracellular Targets. *Nat. Commun.* **2016**, *7* (1), 10778. https://doi.org/10.1038/ncomms10778.
